# Combined Partial-Nitrification and Phosphorus Removal with the co-Existence of Nitrite-resistant phosphorous accumulating organisms (PAOs) and nitrifiers in the treatment of high-strength manure digestate

**DOI:** 10.1101/2023.11.29.569322

**Authors:** Yuan Yan, Peibo Guo, Mathew Baldwin, Guangyu Li, Hyun Yoon, Philip McGuire, Yi Sang, Matthew C. Reid, Joseph Rudek, April Z. Gu

## Abstract

Concurrent biological phosphorus (P) recovery and nitrogen (N) removal in treating high-strength wastewater (such as anaerobic digestate) has been considered incompatible due to presumed conflicts in the conflicting optimum conditions required by phosphorous accumulating organisms (PAO) and nitrifiers. However, this study achieved a stable nitrite accumulation while still maintained PAO activities in one sequencing batch reactor for treating the manure digestate under two aeration schemes (continuous versus intermittent aeration). Nitrite accumulated up to 80.5 ± 21.1 mg-N/L under continuous aeration (6 h) mode. Switching to intermittent aeration (equivalent to 3 h) halved nitrite accumulation but increased total nitrogen removal efficiency from 53.5 ± 12.2% to 84.7 ± 9.4%. Mass balance analysis indicates that nearly all ammonia was removed as N_2_O. Both Enhanced Biological Phosphorus Removal (EBPR) activity assessment and phenotypic trait detection via single cell Raman spectrum (SCRS) confirmed the existence of yet to be identified PAOs that are resistant to high nitrite inhibition in our system. Visual Minteq calculation indicates that high concentrations of Ca in manure digestate may form precipitates and influence the bioavailability of P forms. Therefore, both biotic and abiotic pathways lead to a total P removal rate around 61.0 ± 6.8%. This study highlights new opportunities to combine short-cut nitrogen removal via partial nitrification, nitrous oxide (N_2_O) collection, and EBPR in commercial farm-collected digested manure wastewater. Higher N and P removal efficiency could potentially be achieved by tuning aeration schemes in combination with down-stream anammox process.

**Synopsis:** Concurrent partial nitrification, N_2_O accumulation, and EBPR activity were found, leading to the exploration of novel nitrite-resistant PAOs, simultaneously N/P recovery, and waste-energy conversion in treating high strength wastewater.

## 1. Introduction

In recent years, there has been a growing interest in developing manure management technologies that focus on resource recovery of water, biosolids, struvite, biogas, and bioplastics.^1^ Digested manure comprises relatively high concentration of soluble chemical oxygen demand (sCOD) ranging from 700 to 2500 mg/L, ammonium as 47-1300 mg N/L, phosphate as 10-120 mg P/L.^2^ For farms with sufficient land, energy recovery can be realized through the capture of biogas generated via anaerobic digestion of manure wastewaters containing high-strength organic matter, while the remaining nutrient, mainly nitrogen and phosphorus, could be used as fertilizer. However, many concentrated animal feeding operations (CAFOs), which contain more than 1,000 cows per dairy farm, have limited land area and therefore cannot accommodate the nutrients produced by digestion of their manure.^3,4^ If all digested manure were land applied, N and P in excess of crop needs could be transported into waterbodies as runoff, leaching and erosion and contribute to aquatic ecosystem eutrophication.^5^

Since manure is a byproduct of livestock production, farm owners are responsible for managing manure that is produced on their farms. An effective technology with less chemical addition and lower operation cost is crucial for the operation of a sustainable land-limited CAFO. In this context, improving the biological nitrogen (N) and phosphorus (P) removal in the current manure management chain is desirable. However, the environmentally-sound enhanced biological nutrient removal technology, widely practiced for municipal wastewater, has hardly been explored for high-strength agricultural or food wastewater.^6^ A few studies on biological treatment of manure digestate have been reported, but they primarily focused on either N or P removal alone.^7–10^ Considering the presence of both P and N in these high-strength wastes, the integration of Enhanced Biological Phosphorus Removal (EBPR) with carbon- and energy-efficient short-cut N removal (versus conventional carbon-dependent denitrification) would increase the economic feasibility. Furthermore, unintended greenhouse gas (GHGs) (particularly nitrous oxide - N_2_O) emission from biological nitrogen removal processes has been a concern.^11^ New perspectives have emerged recently suggesting that N_2_O could be used as a fuel additive to enhance the energy generated from methane captured in enclosed anaerobic digester systems.^12,13^ A previous study indicated that 65∼85% influent N could be recovered as N_2_O in the Coupled Aerobic−Anoxic Nitrous Decomposition Operation (CANDO) system.^14–16^ In additional research, high strength digestate was found to be suitable for CANDO systems to recover N_2_O.^13^ Integrating N_2_O recovery with simultaneous N and P removal would greatly enhance the economic feasibility of manure digestate management and increase the possibility to turn the N_2_O side-product, from a waste and climate pollutant into energy.

Though the integration is considered challenging due to presumed conflicts in the design principles and conditions required by these two distinct bioprocesses,^17,18^ our research has achieved simultaneous N and P removal (> 97%) with stable nitrite accumulation in a lab-scale sequencing batch reactor (SBR) treating synthetic manure wastewater with NH_4_^+^-N 50-100 mg-N/L in a prior study.^19^ This result gave us confidence that if the Side Stream EBPR (S2EBPR) concept were extended to anaerobic phases to encourage hydrolysis and fermentation of sludge incorporated in the system, it could be combined with the short-cut N removal process to achieve simultaneous N and P removal from high strength wastewater. Nevertheless, the feasibility of transitioning this configuration to dairy manure digestate collected from a commercial dairy is still challenging due to several factors. Firstly, the characteristics of manure digestate vary depending on its origin (swine, cattle, hog) and the digestion parameters (summarized in Table S1) pose difficulties in directly applying optimized operation parameters from one type of manure digestate to another.^20^ In addition, the presence of high metal concentrations and organic forms of P in commercial manure digestate may affect the bioavailability of ortho-P that PAOs required.^21–23^ Furthermore, manure digestate is typically characterized as having high soluble Chemical Oxygen demand (sCOD)/P ratio (10-30) but low volatile fatty acids (VFAs) concentration.^24–27^ This particular nutrient composition poses limitations on the growth of PAO.^28–30^

In this study, we further explore the feasibility of simultaneously coupled EBPR with shortcut N removal for treating commercial manure digestate wastewater. In a lab-scale SBR, both N and P removal performance were monitored by routine chemical analyses of influent and effluent. Both P and N removal activities, including stoichiometry and kinetics, were evaluated via batch activity tests. The greenhouse gas emissions were monitored during the SBR operation cycles of the reactor. This research will advance our understanding of combining short-cut nitrogen removal with S2EBPR in treating manure digestate.

## 2. Materials and Methods

### 2.1 Commercial manure digestate

The manure slurry was obtained from a collection pit next to the two anaerobic digesters in a dairy CAFO near Ithaca, New York, and was stored at 4°C in polyethylene jugs until use. Due to the high solid content and nutrient concentration in the raw manure slurry, the manure slurry was centrifuged, and the supernatant was diluted 10 times prior to feeding into the reactor to prevent bio-inhibition effect induced by extremely high ammonia concentration. The composition and concentration of manure are listed in Table 1, being similar to the manure digestate wastewater reported previously (Table S1).

**Table 1.** Manure digestate components in influent (10X diluted)

| Parameter | 2020-Period-I | 2021-Period-II |
| --- | --- | --- |
| pH | 7.51 | 7.5-7.88 |
| sCOD (mg/L) | 959-2200 | 1379-1922 |
| BOD<sub>5</sub> (mg/L) | 558-657 | 963 |
| TOC (mg C/L) | 224-270 |  |
| TP (mg P/L) | 36-50 | 20-33 |
| PO<sub>4</sub> (mg P/L) | 10-35 | 4-11 |
| NH<sub>4</sub> (mg N/L) | 182-230 | 180-264 |
| TSS (mg/L) | Both ranged between 2,290-3,760 |  |
| VSS (mg/L) | Both ranged between 1,740-2,379 |  |
| VFA (mg C/L) | 153 | 250 |
| C/N (mg sCOD/mg NH<sub>4</sub><sup>+</sup>-N) | 5.6-8 | 7.2-7.9 |
| C/P (mg sCOD/mg PO<sub>4</sub><sup>3-</sup>-P) | 54-60 | 150-174 |

### 2.2 SBR operation

A lab-scale SBR with a working volume of 4 L and a volumetric exchange ratio of 50% was used in this study. The reactor was inoculated with EBPR activated sludge from Hampton Roads Sanitation District (HRSD) Virginia Initiative Treatment Plant (Norfolk, VA) and Nansemond Treatment Plant (Suffolk, VA).

The operation periods were separated into two modes. In Period-I (97 days), the reactor was operated for anaerobic/continuous aerobic phases as follows: Feeding time for 5 min, anaerobic phase for 2.5 h, aerobic for 5 h, settling time for 20 min, decanting, and idle for 10 min. In Period-II (87 days), the reactor was operated for anaerobic/intermittent aerobic phases as follows: Feeding time for 5 min, anaerobic phase for 2.0 h, 30 min on/off aeration for 5 h, settling time for 20 min, decanting, and idle for 10 min.

During the operation, pH was not controlled and varied between 7.2-8.5. Dissolved oxygen (DO) was between 2-8 mg/L during the aerobic phase to avoid any oxygen limitations (Figure S1). The reactor was operated at a constant temperature of around 25 °C. The sludge retention time (SRT) was kept at 10 days by periodic wasting of the mixed liquor at the end of the aerobic phase and the volatile suspended solids (MLVSS) was maintained around 6600 mg/L.

### 2.3 Performance monitoring and chemical analysis

The influent and effluent samples of the reactor were sampled routinely and then centrifuged immediately at 6000 g for 10 min. The supernatant was filtered by 0.22 μm filters before chemical analysis. The untreated samples were stored at -20 °C for further analysis. The concentrations of NH_4_ ^+^, NO_3_ ^−^, NO_2_ ^−^, and PO_4_ ^3−^ were measured by ion chromatography (IC) (Thermo Fisher Scientific, Dionex ICS-2100, USA) with anion and cation exchange columns, respectively. Mixed liquor suspended solids (MLSS), and MLVSS were analyzed in accordance with Standard Methods.^31^ Total and soluble COD (Hach reaction digester method) and the total VFA concentration (Hach esterification method, TNT872, USA) were measured by colorimetric spectrometer.

The pH and DO concentration in the reactor were routinely measured by DO and pH meter kit (DFRobot, Shanghai, China). The pH, DO and the oxidation reduction potential (ORP) in the influent, effluent, and at the end of anaerobic phase were measured by Portable Dedicated pH/ORP/mV Meter (Hach Company, HQ1110, USA).

For Al, Fe, Ca, Mg and total phosphorus (TP) measurements, samples were filtered, digested, and measured by inductively coupled plasma mass spectrometry (ICP-MS) (Agilent 7800, USA). Soluble reactive phosphorus (SRP) was measured through molybdate blue method 4500-P.^31^

### 2.4 Activity batch tests

Denitrification and anammox activity tests were performed to identify the contribution of nitrogen removal pathways. Activated sludge was washed 3 times to remove the remaining medium. After that, about 3 g wet sludge was collected into a 250 ml serum bottle and concentrated medium were injected into bottle to initiate the reaction. After 5-minute premixing, samples were collected every 30 minutes for a total of 2.5 h, filtered through a 0.22 μm filter and then analyzed for NH_4_ ^+^-N, NO_2_ ^−^-N, NO_3_ ^−^-N concentrations. At the end of batch tests, the total suspended solids (TSS) for each bottle were measured according to the standard methods.

The final concentration of media (sodium acetate, nitrate, nitrite, ammonium) in each batch tests, the nitrate reduction, nitrite reduction rates, and total nitrogen removal rates in anammox tests were respective calculated by linear regression and shown in Table S2.

### 2.5 N_2_O measurement

For gaseous N_2_O sampling, the holes on top of reactor for sensors were sealed with rubber stopper. Only one hole was outfitted with a vent for gas collection. A small fan was installed inside of the reactor to achieve thorough mixing of the headspace gas. Gas samples were collected from the sampling vent into 50 ml nylon syringes at specific time intervals. During anoxic phases, it is assumed that gaseous N_2_O increases linearly with microbial activity. Therefore, gas samples were only collected at the end of each anoxic phase. During aeration phases, since dissolved N_2_O would be stripped out of the liquid phase, more intensive sampling points were used so the gaseous N_2_O emissions can be fitted with an exponential equation.

For dissolved N_2_O measurement, 5 ml mix liquor was collected every 30 minutes during the cycle. After shaking 3 minutes with 45 ml N_2_, the equilibrated gas samples were stored in gas vials for quantification.

N_2_O was measured via gas chromatography (GC) analysis (Shimadzu GC-2014, Kyoto, Japan), using an electron capture detector, and CH_4_ was measured with a flame ionization detector. All N_2_O emissions (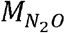, mg-N_2_O-N) were calculated by integrating the gaseous N_2_O curve in aeration phase and anoxic phase using equation (1):

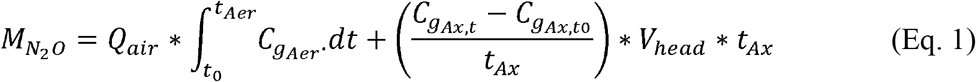

Where:*Q*_air_ is the flow rate of air, 6.7 L/min;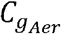 is the concentration of N_2_O in aeration phase, mg-N_2_O-N/L;*t*_*Aer*_ is the aeration time, 30 min in each aeration phase;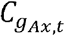 and 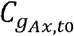 are the concentration of N_2_O at the end/begin of anoxic phases, mg-N_2_O-N /L;*t*_*Ax*_ is the anoxic time, 30 min in each anoxic phase;*V*_*head*_ is the head space of the reactor, 6 L.

N_2_O emission factors were calculated as a percentage of the average influent NH_4_ ^+^-N load being emitted as N_2_O.

### 2.6 Single cell Raman spectrum (SCRS) analysis and phenotypic profiling

Sludge samples (1 ml) were collected from the reactor at the end of the aerobic phases and centrifuged for 5 minutes at 6000 rpm. After being washed three times with 1X Phosphate Buffered Saline (PBS), 50X diluted samples were homogenized by pushing the cells through 26-gauge needle syringe 10 times to obtain uniform distribution of cells and then 2.5 ul samples were prepared on optically polished CaF_2_ windows (Laser Optex, Beijing, China). Raman spectra of single-cell were scanned from 400 to 1800 cm^-1^ by a LabRam HR Evolution Raman microscope equipped with a magnification of x50 objective (HORIBA, Japan). Spectrums were acquired according to previously published methodologies ^32,33^. The phenotyping analysis was conducted according to previously published research.^34^. The spectra data were processed with the smoothing and filtering, background subtraction, baseline correction steps by the software LabSpec 6 (Detailed key parameters were in supporting information Text S1).

The presence of polyhydroxyalkanoates (PHA) in single cells was identified by the signature peaks in the range of 1715-1740 cm^-1^ and the presence of polyphosphate (poly-P) was identified by the two signature peaks in the range of 685-715 cm^-1^ and 1150-1180 cm^-1^ based on previous study ^35^. The relative intensity of PHA and poly-P for single cell was normalized by the intensity of amide I vibration at 1640-1690 cm^-1^. The hierarchical clustering analysis (HCA) was applied on all of the single-cell Raman spectra from activated sludge samples to obtain phenotypic profiles based on operational phenotypic units (OPUs) according to our previous study.^34^ The cosine similarity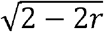, r-correlation efficient) was used to measure metrics between two samples, average linkage was applied to quantify dissimilarities between two clusters and the cutoff threshold for OPUs was set with Akaike information criterion (AIC). The Raman fingerprints of Cte probe ^36^ hybridized cells were used as the *Comamonadaceae*-labeled reference.

### 2.7 DNA extraction and 16S rRNA gene amplicon sequencing

To investigate the microbial community dynamics, sludge samples were collected at the end of the aerobic phase periodically and the genomic DNA was extracted using the FastDNA Spin Kit for Soil (MP Biomedical, USA).. The extracted DNA was sent to University of Connecticut-MARS facility for PCR amplification and sequencing targeting the V4 region using the primers 515F (5’-GTGCCAGCMGCCGCGGTAA-3’) and 806R (5’-GGACTACHVGG GTWTCTAAT-3’) and the amplicons were sequenced on the Illumina MiSeq with V2 chemistry using paired-end (2 x 250) sequencing. The raw paired-end reads were assembled for each sample and processed in Mothur v. 1.36.1 following the MiSeq standard operating procedure (SOP).^37^ High-quality reads were obtained after quality control and chimera screening and then clustered at a 97% similarity to obtain the operational taxonomic units (OTUs).

### 2.8 Process Modeling

The SUMO™ 19 process modeling package by Dynamita (Nyons, France) was used to evaluate the SBR operation. The SUMO2 model utilizing two-step nitrification and denitrification, and the Barker-Dold model for P removal was selected. Chemical precipitation was enabled along with pH and DO calculation.

Analysis of phosphorus speciation was further conducted using Visual MINTEQ version 3.1.^38^ For speciation, the input data were concentration of the NH_4_ ^+^-N and NO_2_ ^−^-N, Ca^2+^, Mg^2+^, pH value, ORP, and DOC at the beginning of reaction, the end of anaerobic stage and in the effluent (Table S3). For assessing the ion-binding behavior of humic acids, the NICA-Donnan model was employed with default parameters. This model assumes a continuous distribution of deprotonated carboxylic and phenolic groups and was widely applied in complex wastewater.^39^

## 3. Results and Discussion

### 3.1 Reactor performance

#### 3.1.1 Period-I (anaerobic/continuous aeration)

In Period-I, the SBR was operated for 97 days with DO concentration in aerobic phase varied from 2.0 to 6.0 mg/L. The effluent NH_4_^+^-N gradually decreased to zero with a stable nitrite accumulation after day 43. The effluent NO_2_^−^-N concentration increased to 80.5 ± 21.1 mg-N/L with the effluent NO_3_^−^-N of 0-3 mg-N/L (Figure 1). The nitrite accumulation rate (NAR), calculated as the ratio of nitrite to nitrite and nitrate, could reach 97.2 ± 2.1%. The overall total nitrogen removal efficiency (TNRE) is around 53.5 ± 12.2%.

Initially, the PO_4_ ^3−^-P removal rate could reach 60% upon seeding with EBPR sludge. However, the P removal rate gradually decreased to 20% when the system began to accumulate nitrite. Though a stable P and N removal could be achieved with 20.4 ± 6.4 mg N/L nitrite accumulation in a similar system,^19^ the present study encountered a higher nitrite accumulation of nearly 80 ± 21 mg-N/L, which could greatly hamper the activity of both N removal and P accumulating bacteria. To address this issue, the continuous aerobic phase (6 h) was changed into multiple 30/30 mins aerobic/anoxic phases in Period-II to alleviate the possible nitrite inhibition effect.

**Figure 1.**
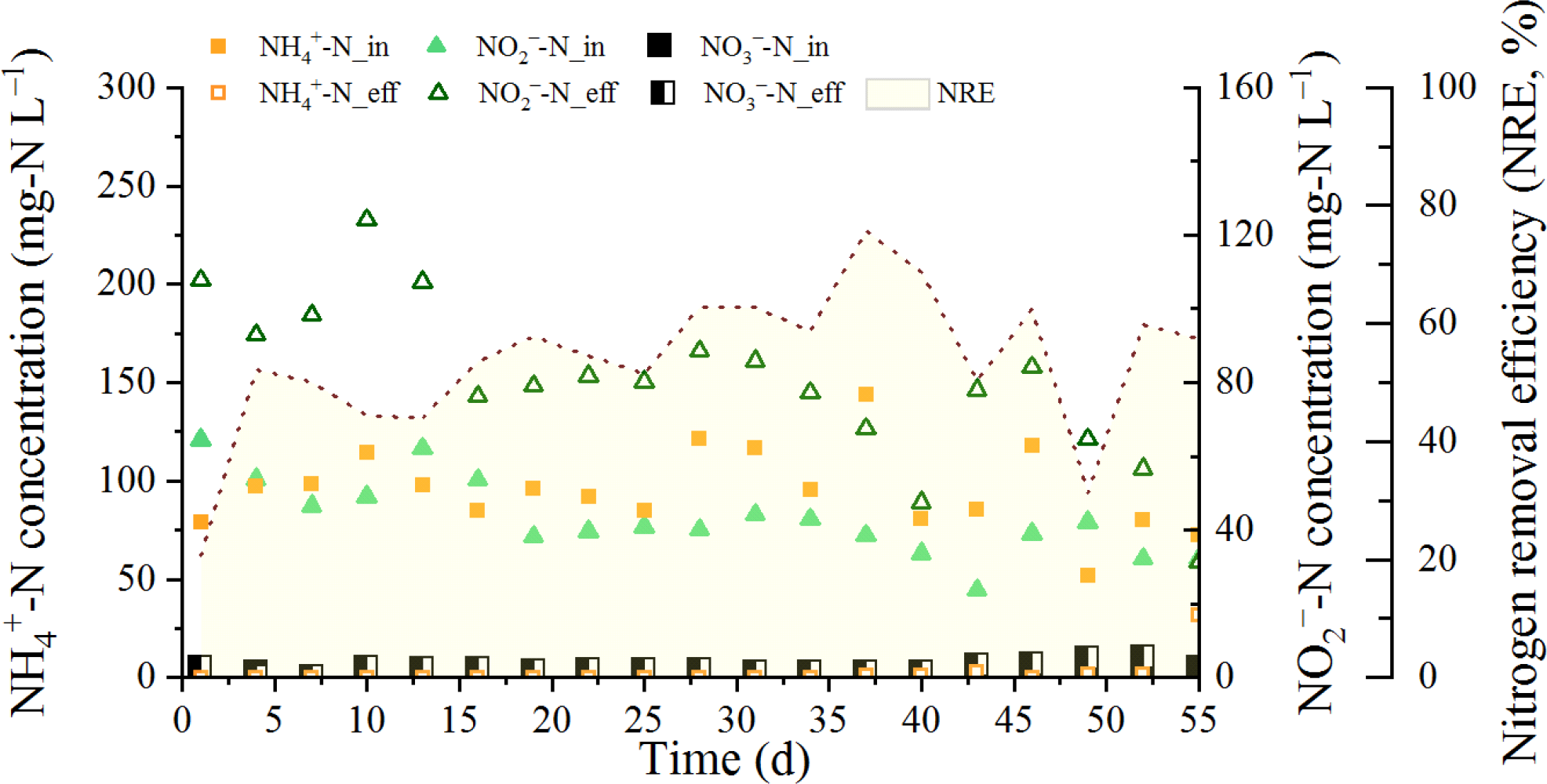
Reactor performance in Period-I, dates are readjusted to 1 when the system reached steady state.

#### 3.1.2 Period-II (anaerobic/Intermittent Ax/Aer)

In Period-II, the influent ammonia concentration was maintained the same as in Period-I. Once the reactor reached steady operation, it was maintained for 90 days. Reducing the aeration time to 3 hours resulted in an increase of effluent ammonia to 16 ± 9.5 mg N/L, while the effluent nitrite reduced to 27.6 ± 9.6 mg N/L (Figure 2). There was no detectable nitrate in Period-II with NAR maintained at 100%. The overall TNRE increased to 84.7 ± 9.4%, indicating a stronger denitrification activity facilitated by a prolonged anoxic phase (3 h). The PO_4_^3−^-P removal rate seems higher than in Period-I but that was due to the decrease concentration of soluble P in the raw manure digestate (Table 1). The actual SRP reduction still maintained around 3-5 mg-P/L per cycle, indicating that P removal performance was not influenced by either lower soluble P or the reduced nitrite accumulation in Period-II relative to Period-I.

**Figure 2.**
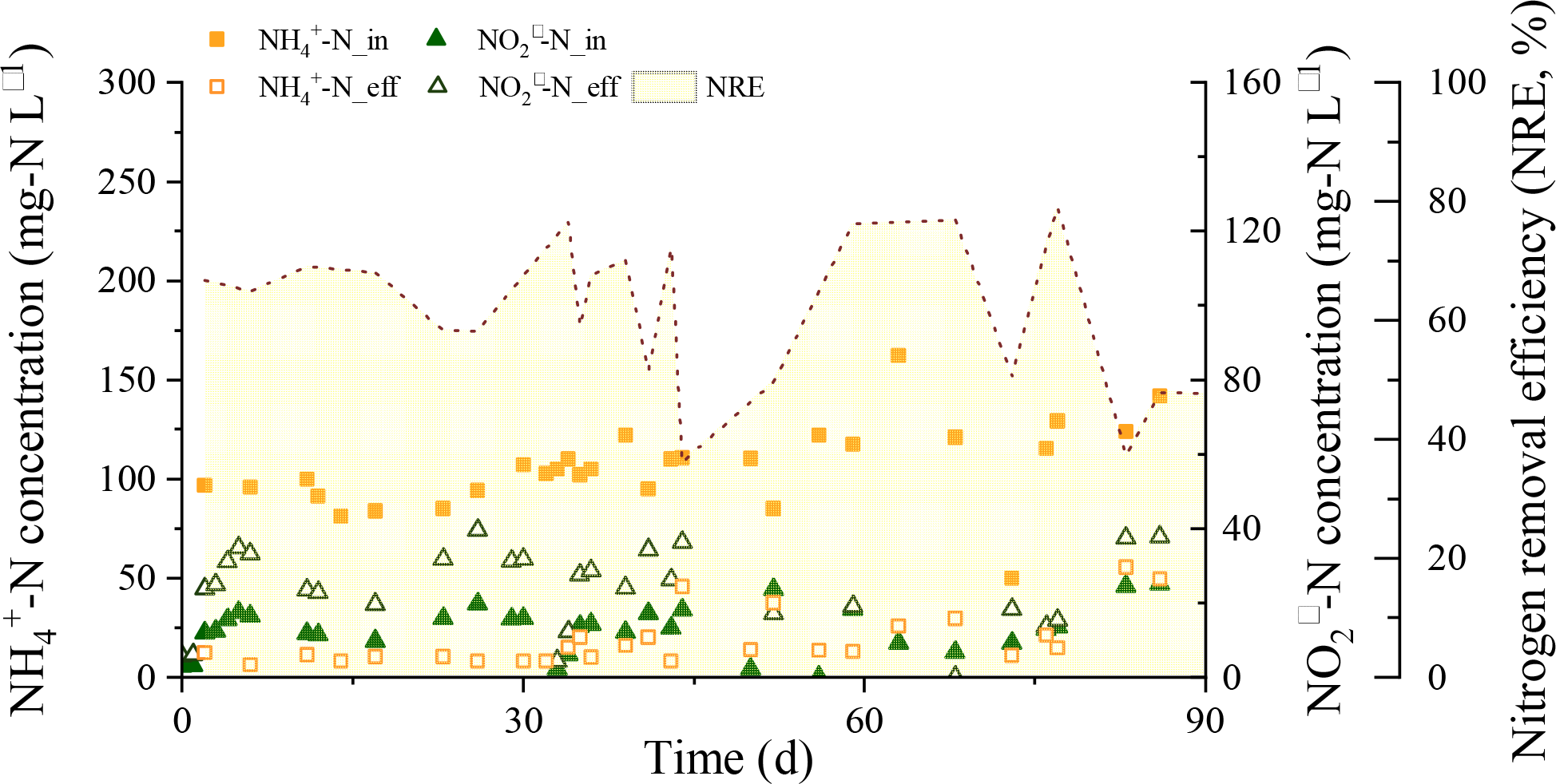
Reactor performance in Period-II.

### 3.2 Stable nitrite and N_2_O accumulation during an SBR operation cycle in Period-II

#### 3.2.1 Nitrite accumulation due to partial nitrification

Though the average DO concentration (∼6 mg/L) in aeration phases are much higher than the setpoint in conventional partial nitrification reactor (0.5∼2 mg/L), a constant nitrite accumulation was observed under two periods. SBR operation cycles were analyzed on day 11, 23, 25, 30, and 40 in Period-II. The residual nitrite from the last aeration stage was fully denitrified in the subsequent 2-h anaerobic phase (Figure 3). DO concentrations in the first three aerobic stages varied from 1.0 to 3.0 mg/L, being similar to the conventional partial nitrification reactor. Subsequently, the DO concentration increased gradually to 8 mg/L during the following three aerobic stages. Around10-24 mg-N/L NH_4_ ^+^ was oxidized into NO_2_^−^ with no nitrate formation throughout the cycle. These results indicated a strong Nitrite Oxidizing Bacteria (NOB) inhibition.

**Figure 3.**
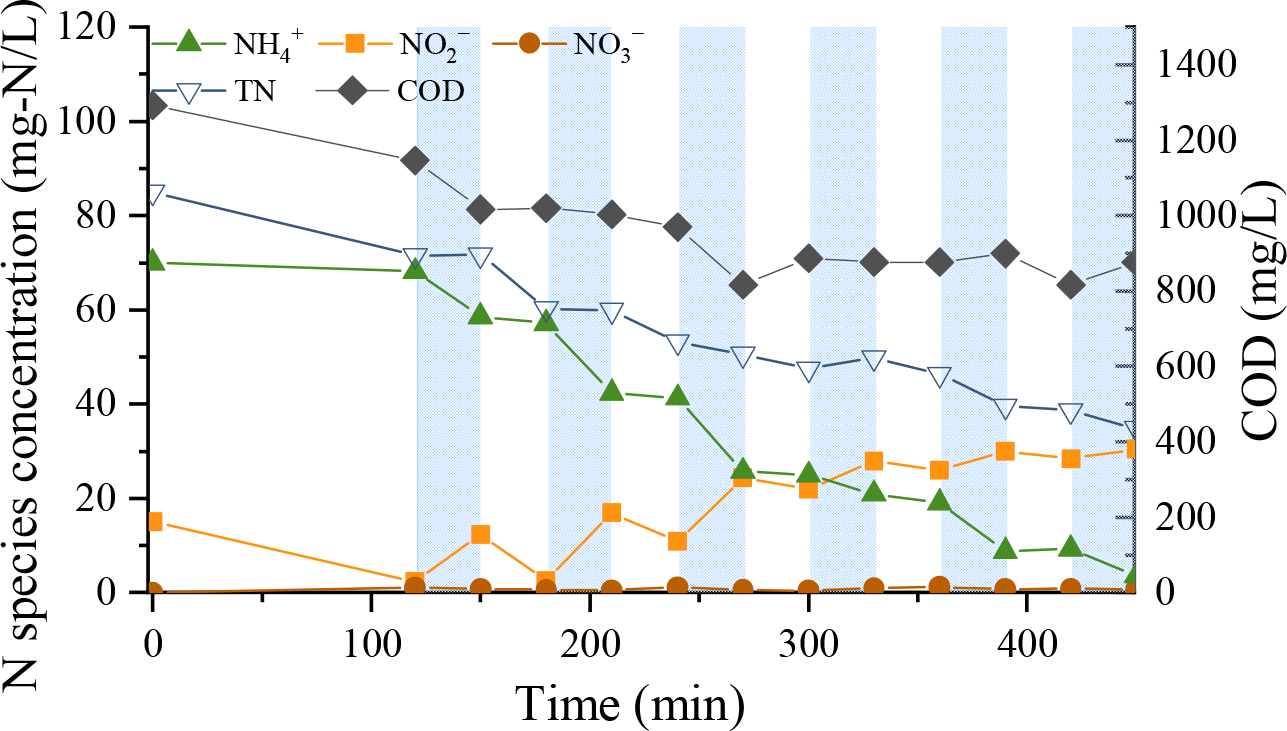
sCOD and nitrogen species in a typical cycle in Period-II (day 40)

In the first several anoxic phases, the NO_2_ ^−^ reduction could reach∼8 mg/L per half hour. As the sCOD no longer decreased in the final two anoxic phases, only ∼2 mg/L NO_2_ ^−^ could be reduced. It’s also interesting to find 0.7-1.8 NH_4_ ^+^-N/L disappeared along with nitrite reduction in anoxic phases, which implies the existence of denitrification and/or anammox activity.

To further identify the N loss during the aerobic and anoxic phases, we conducted serial batch tests (Table S2). With external acetate addition, nitrate reduction rate was 2.94 ± 0.09 mg-N/g-VSS/h with no nitrite accumulation. Similarly, with acetate and NO_2_^−^-N as the sole electron acceptor, the NO_2_^−^-N reduction rate is 2.77 ± 0.13 mg-N/g-VSS/h, being comparable to that of NO_3_ ^−^-N reduction rate. If acetate was not present in the test media, the denitrification rate is around 0.20 ± 0.01 mg-N/g-VSS/h, being similar to the normal endogenous denitrification rate in the range of 0.2−0.6 mg N/h g MLVSS. ^40^

With ammonia, nitrite and no organic carbon source, we obtained anammox activity as 1.32 ± 0.03 mg-N/g-VSS/h. If the denitrification and anammox activities were both active in the first two anoxic phases, then ∼4100 mg/L VSS in the reactor could remove ∼8.1 mg/L NO_2_ ^−^-N, in accordance with the anoxic N loss obtained in SBR cycle analysis (Figure 3). When bacteria consume all bioavailable carbon in the first several Aerobic/Anoxic (Aer/Ax) phases, heterotrophic denitrification rate ceased and only leave anammox bacteria for nitrogen removal. Based on calculation, anammox activity alone will contribute to ∼2.6 mg/L TN removal that are similar to the last three anoxic phases (Figure 3). The occurrence of stable partial nitrification in both periods offers a promising opportunity for integrating a downstream anammox process into manure management practices. This integration could serve as an effective strategy to further reduce effluent TN levels.

#### 3.2.2 N_2_O accumulation profile over the course of a SBR operation cycle

In addition to concerns regarding eutrophication resulting from manure wastewater overapplication to croplands, the accumulation of nitrite and the aeration process also raise concerns about greenhouse gas emissions. According to field sampling report for full-scale domestic waste water treatment plants (WWTPs), the N_2_O emission factor varies largely from 0.0001 to 0.112 kg N_2_O-N/kg NH_4_ ^+^-N_influent_, depending on the reactor configuration and process characteristics and measurement methodologies in a set of 29 WWTPs all over the world.^41^ The N_2_O emissions from manure digestate has been found to generally be higher than from municipal wastewater, ranging from 8.2-11% of removed NH_4_^+^-N.^8,42^ With synthetic high strength wastewater (1700-1800 mg/L NH_4_ ^+^-N_inf_), 20–30% of influent nitrogen was emitted as N_2_O.^43^ Previous researchers obtained > 90% N_2_O emission when they feed nitrite in a simultaneous nitrification, denitrification, and phosphorus removal SBR. In these studies, N_2_O was mainly produced under high nitrite and high DO situations.^44^ The characteristics of swine waste was also proven to inherently influence the activity of NOB and/or N_2_O production during ammonium oxidation.^8^

Since the solubility of N_2_O is relatively high, the N_2_O generated in denitrification reactor may dissolve in solution and be stripped out by the following aeration. To evaluate the gas emission in our biological reactor, N_2_O, CO_2_ and CH_4_ concentrations in both dissolved liquid and headspace of our reactor were analyzed during a SBR cycle (Figure 4). At the end of anaerobic stage, only CH_4_ accumulated in liquid phase, and it was stripped out within 2 aeration periods (Figure 4A). N_2_O concentration at the end of each anoxic phase varies from 5.8-9 mg-N/L. N_2_O concentrations at the end of aerobic phases are much lower than anoxic phases, maintaining around 1 mg-N/L. Similar variations were observed for N_2_O and CH_4_ concentrations in the headspace, although they were consistently 1-2 orders of magnitude lower than the dissolved concentrations. Intensive sampling in aeration phase (Aeration phase 4) confirmed that the N_2_O concentrations in headspace peaked when aeration was on, and subsequently decreased exponentially due to air stripping (Figure S2). The total emitted N_2_O can be calculated by Equation 1.

**Figure 4.**
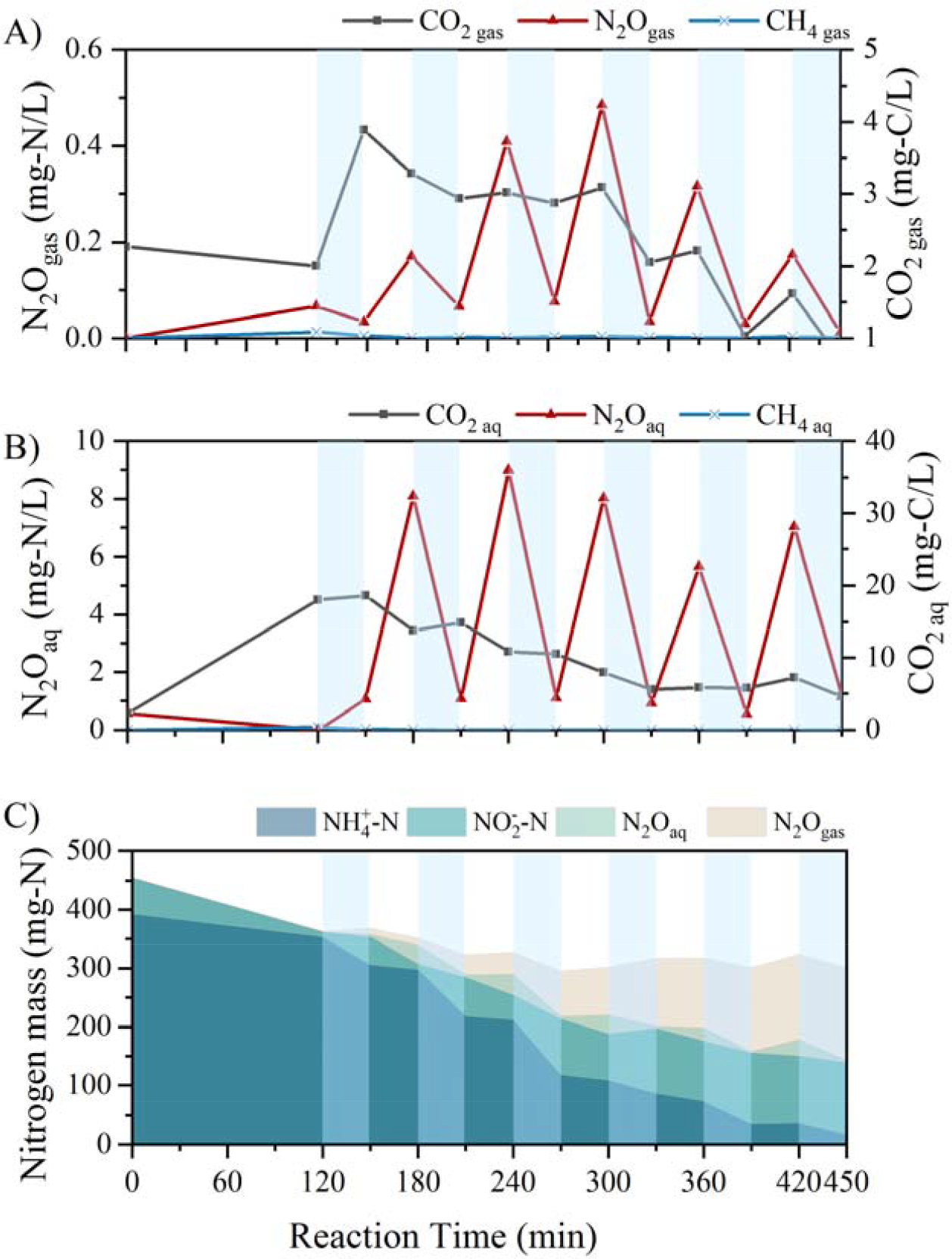
(A) Gaseous N_2_O, CO_2_, and CH_4_; B) Dissolved N_2_O, CO_2_, and CH_4_; and C) Nitrogen species mass variations in a typical cycle in Period-II (day 85).

Taking both aqueous and gaseous nitrogen species into account, the nitrogen conversion during a SBR cycle is summarized in Figure 4C. During the anoxic stages, N_2_O accumulated in aqueous phase. Subsequent aeration led to the stripping of dissolved N_2_O into the gaseous phase. It is worth noting that the decrease in the mass of dissolved N_2_O corresponds closely to the increase observed in the gaseous phase. This observation strongly suggests that the primary source of N_2_O is the anoxic phase rather than the aerobic phases. The incomplete denitrification pathway, specifically the conversion of NO_2_^−^ to N_2_O, significantly contributes to the presence of N_2_O in our system. The N_2_O emission factor in total cycle is as high as 0.4 kg N_2_O-N/kg NH_4_^+^-N_influent_. Our findings aligns with the CANDO system’s approach of using denitrifiers to accumulate N_2_O as end products from NO_2_^−^-rich wastewater.^12,13^

### 3.3 Interference of Abiotic Phosphorus Precipitation with Biotic Phosphorus Removal

In the presence of PAO activity, intracellular poly-P would be hydrolyzed into orthophosphate in the anaerobic phase to provide energy for Polyhydroxyalkanoates (PHA) synthesis, leading to a notable SRP increase in EBPR system. However, during a typical Anaerobic/Aer/Ax cycle, we only observed 2.08 mg-P/L release at the end of anaerobic phase (Figure 5). Initially, we attributed this limited release to the inhibition caused by high nitrite concentrations. However, when we switched to measuring soluble total phosphorus (STP) using ICP-MS, the results clearly demonstrated a total phosphorus release of 9.6 mg-P/L after the 2-hour anaerobic phase. The discrepancy between STP and SRP measurements suggests that 6.92 mg-P/L was released in the form of other P species that cannot be accurately quantified as SRP using molybdate-based methods.

**Figure 5.**
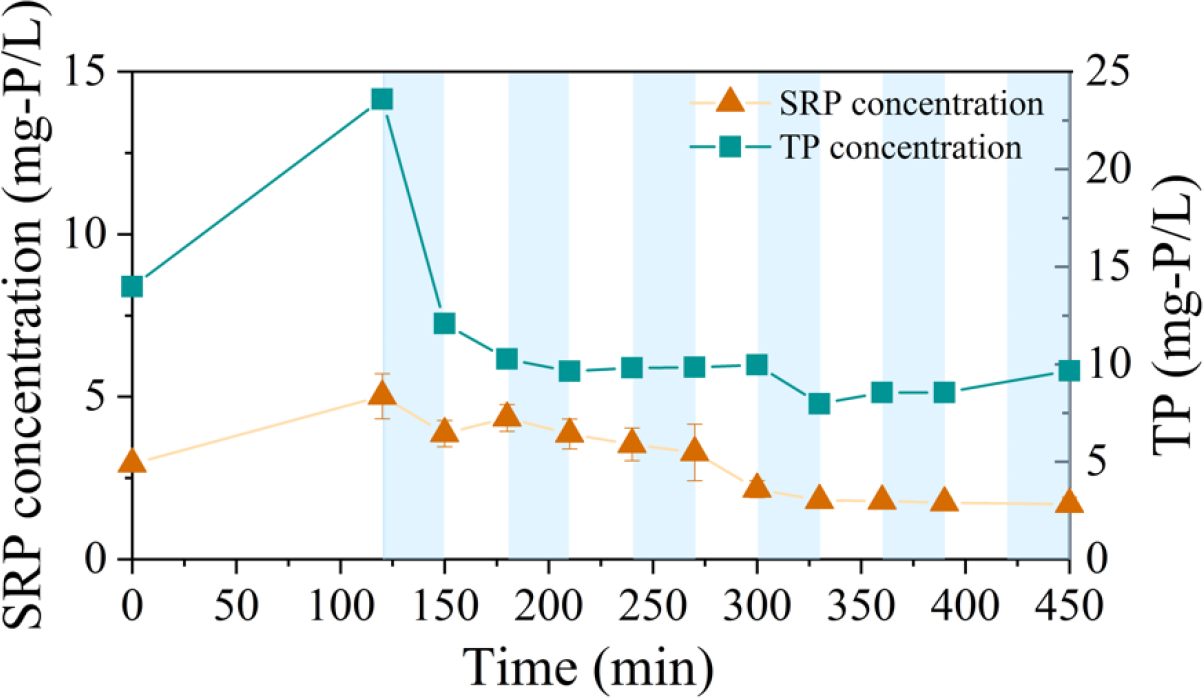
SRP variations in a typical cycle (Period-II, day 85)

Our manure digestate influent contained significant concentrations of Ca^2+^ and Mg^2+^ ions, measuring 68.2 ± 28.8 mg/L and 40.0 ± 16.5 mg/L, respectively. These elevated levels of cations have the potential to influence the metabolic pathways of PAOs^45^ and can also lead to the formation of precipitates when combined with released SRP, thereby affecting PAO activity.

SUMO process model adopted the kinetic rate expression to integrate the chemical–physical processes in the same reactor.^46^ It was predicted that the metastable precipitation (i.e., amorphous calcium phosphate, ACP) would be stable in wastewater instead of transforming to thermodynamically stable species, such as hydroxyapatite. Enabling precipitation calculation, SUMO process model predicts the occurrence of ACP in reactor would be approximately 1100 mg/L due to the high Ca^2+^ concentration, representing 12% of the TSS. However, this predicted amount of ACP does not align with our observations. This calculation fails to capture the biological phosphorus release and uptake, as depicted in Figure S3. The precipitation processes in wastewater are also influenced by the presence of inorganic and organic ligands. It should be noted that the anaerobic digestion process would lead to an increased proportion of humic-like substances in manure digestate.^47^ The distinct composition of digestate compared to municipal wastewater could have influenced the speciation outcomes in the SUMO model. The elevated proportion of organic matter may have contributed to the stabilization of fine particles within the colloidal size range as indicated previouly.^48^ Furthermore, the presence of humic acid can also impact the cationic bridging facilitated by divalent cations such as Ca^2+^ and/or Mg^2+^. This can lead to the formation of bidentate or ternary complexes between phosphate and dissolved organic matter.^49^

To gain a better understanding of the influence of organic ligands, we use NICA-Donnan model to calculate P precipitation at different stages of the cycle, including the beginning of the cycle (T0), the end of the anaerobic phase (T120), and the effluent (Eff.) with changes in NH_4_ ^+^-N, NO_2_ ^−^-N, pH, ORP, VFA, TP, SO_4_ ^2−^, Cl^−^, Ca, Al, Mg, and Fe (Table S3). We assume that newly the formed ACP and/or humic acid (HA)-Ca-P complexes are in colloidal size fraction and therefore pass through 0.45 um filter. In this context, only SRP is readily bioavailable. If all organic P in influent is sufficiently hydrolyzed into orthophosphate, then the model would predict 12.1 mg-P/L precipitation as hydroxyapatite (**Table 2**). After 120 min anaerobic phase, TP concentration increased 9.6 mg-P/L, indicating an anaerobic P release activity by PAOs. Accounting the released PO_4_ ^3-^ (Figure 5), P precipitation was modelled to increase to 18.82 mg-P/L (**Table 2**). The remaining 4.8 mg/L dissolved PO_4_ ^3-^ is in accordance with the measured SRP at the end of anaerobic phase (5.0 mg-P/L in Figure 5). Towards the end of the reaction, ammonia was fully oxidized to nitrite accompanied by a decrease in pH to 6.6, leading to a redissolution of phosphorus precipitates. However, the released phosphate was not captured by either SRP measurement or the STP measurement. It may have been stored intracellularly by bacteria. The precipitation/dissolution of hydroxyapatite in anaerobic/aerobic cycle has also been reported by Zhang et al., (2015).^45^ With precipitation model, the phosphorus release/uptake was calculated as 8.8 and 12.4 mg-P/L, being close to the measured TP release/uptake, indicating that chemical precipitation interferes with the biological phosphorus removal.

**Table 2.** Phosphorus precipitation calculated by Visual Minteq (mg-P/L)

| Time | Measured $\text{PO}_4^{3-}$ (SRP) | Obs. SRP Release/uptake | Measured TP | Obs. TP Release/uptake | Model precipitate | Model Release/Uptake |
| --- | --- | --- | --- | --- | --- | --- |
| $T_0$ | 2.9 | | 14.0 | | 12.1 | |
| $T_{120}$ | 5.0 | +2.1 | 23.6 | +9.6 | 18.8 | +8.8 |
| Eff | 1.7 | -3.3 | 9.7 | -13.9 | 9.8 | -12.4 |

Therefore, the TP release/uptake was 9.6 and 13.9 mg-P/L, respectively (**Table 2**), resulting an P release and uptake rates of 1.1 and 1.05 mg-P/g-VSS/h. This rate is close to the rates obtained in S2EBPR facilities.^18,50^

### 3.4 Nitrite-resistant PAOs and Putative N_2_O generator

Based on 16S sequencing, *Nitrosomonas* was the only known ammonia oxidizing bacteria (AOB) genus that could be detected (0.2%-3.2%) and NOB relative abundance was below the detection limit (<0.1%) (Figure 6), which was similar to the results with synthetic manure digestate.^19^ These results indicate that partial nitrification in high-strength manure digestate is achieved mainly by successful NOB out-selection.

**Figure 6.**
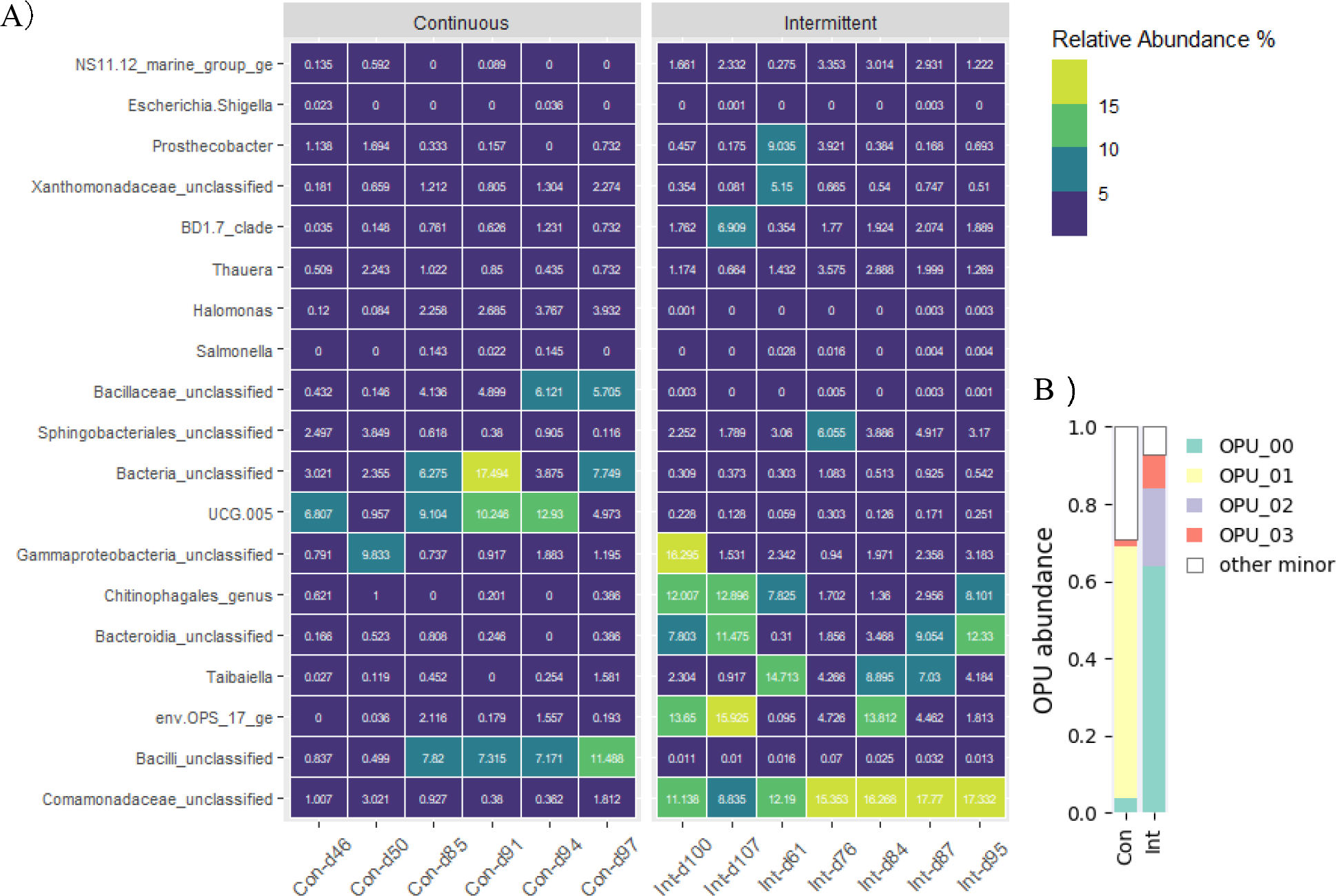
(A) Time series heatmap of the relative abundances of the bacteria in Period-I (Con-) and Period-II (Int-) at the genus level; (B) relative abundance of each operational phenotypic units (OPUs) clustered by the hierarchical clustering analysis (HCA) in Period-I (con.) and Period-II (Int); Values represent average relative abundances (%)

In the synthetic manure digestate reactor, Glycogen Accumulating Organisms (GAOs; *Defluviicoccus* and *Candidatus* Competibacter) (15.6-40.3%) and PAOs (*Candidatus* Accumulibacter) (14.2-33.1%) were the most abundant genus and we postulated that those nitrite-resist PAO members in *Candidatus* Accumulibacter could perform efficient EBPR activities.^19^ However, the microbial communities fed with farm-sourced manure digestate are completely different (Figure 6). In the present study, *Candidatus* Accumulibacter cannot be detected and the only detected GAO, *Candidatus* Competibacter, was below 0.1% throughout the operation periods. In contrast, two putative PAOs,^51^ *Dechloromonas* and *Gemmatimonas* emerged in Period-II (Intermittent aeration scheme) at abundances ranging from 0.1% to 0.6%, which may contribute to the P release/uptake activities in our reactor. It is important to note that 16S sequencing technology can only identify the known PAOs. There is a possibility that there are other bacterial species capable of performing phosphorus uptake and release activities that have not yet been identified. Previous study has demonstrated that Raman spectrum can be employed to differentiate possible PAOs with other Ordinary Heterotrophic Organisms (OHOs) by identify their intracellular poly-P signals at single cell level.^33,35,32^ In the present study, single-cell Raman spectra (SCRS) analysis revealed that approximately 20∼30% cells contain poly-P and ∼10% cells contain PHA at the end of aeration phase in both Period-I and Period-II (Figure S4A and S4B). The high abundance of PHA/poly-P containing cells by Raman detection and low abundance of known PAOs detected by16S amplicon sequencing suggested that some unknown nitrite-resistant PAO could survive with 80.5 ± 21.1 mg-N/L nitrite accumulation and still perform EPBR activity.

The most abundant group shifted from UCF-005 and Bacilli_unclassified (5%-12.9%, 7.2%-11.5%) to Comamonadaceae_unclassified (8.8%-16.6%), env.OPS_17_ge (3%-15.9%), *Taibaiella* (4.3%-14%), Chitinophagales_genus (1.7%-13%) when aeration scheme was changed to intermittent (Period-II). The less common order Chitinophagales has positive correlation and may serve as bioindicators of high P content but low phosphatase activity in soil.^52^ *Taibaiella* is an group of strictly aerobic bacteria that can reduce nitrate to nitrite when oxygen is gone.^53^ Members in the Family *Comamonadaceae* are denitrifiers that are often detected in EBPR systems,^54^ and some of them can degrade solid phase PHA.^55,56^ Since the internal PHA accumulation is the key factor to ensure the stable N_2_O accumulation,^13,16^ the PHA containing cells in our system become the mutual focus of putative nitrite-resistant PAOs and denitrifiers.

All SCRS under two operation periods were gathered to identify the main OPUs based on their phenotypic features, following a developed methodology.^34^ Only one dominant OPU (OPU1) was detected with relative abundance of 11.6% in Period-II (Figure 6B) Corresponding heatmap could be found in Figure S5A. Given that the genus Comamonadaceae_unclassified became dominant in Period-II (Figure 6A), we ran another round OPU analysis by combining all SCRS from present study with the previously obtained SCRS that were labelled with *Comamonadaceae* fluorescent probe. Results showed that OPU1 in the present study is close to the *Comamonadaceae* labelled SCRS (Figure S5B). Notably, all the spectra within OPU1 were featured with poly-P signals in our algorithms, suggesting a potential involvement of these organisms in both P removal and N_2_O accumulation. However, additional studies are needed to confirm phenotype conclusively.

### 3.5 Simultaneous N and P Removal: A promising Application in Manure Management

In the N mass balance of *in situ* batch activity tests, around 55% to 81% N was removed. Remarkably, when calculating the emitted N_2_O by summing up all N_2_O present in both mixed liquor and headspace, it was found that a significant amount of all nitrogen was removed through N_2_O formation (Table 3). While N_2_O has traditionally been considered as hazard intermediates in biological nitrogen removal process due to its high global warming potential, it can also be intentionally stimulated and recovered under well-controlled conditions, like CANDO system, as an oxidant for combustion of biogas methane.^12,13^ The formation of N_2_O during the reduction of NO_2_ ^−^ and oxidation of NH_2_OH is influenced by the presence of nitrite. In our system, a steady accumulation of nitrite was achieved, ranging from 27.6 ± 9.6 mg N/L to 80.5 ± 21.1 mg-N/L (Figure 1 and Figure 2), which is similar to the reported level in coupled aerobic-anoxic nitrous decomposition operation (CANDO). Such high concentration of nitrite accumulation and the switching between anaerobic/anoxic/aerobic condition may contribute to the high N_2_O emission in this study. Considering that manure management often produce biogas through anaerobic digestion,^8^ it is worth exploring the potential to capture N_2_O in the subsequent nitrogen removal process and utilize it as the oxidant in biogas combustion. Though integrating N_2_O as an oxidant may not align with the processing of biogas to biomethane and its subsequent injection into natural gas pipelines, this approach may be practical for on-farm management of biogas collection on large dairies and hog operations with digestate treatment. In addition, it’s crucial to carefully assess the risk of leakage associated with on-farm combustion of a biogas/N_2_O mixture for heat and power generation. Nevertheless, harnessing N_2_O as an energy source could be an advantageous alternative approach rather than striving to minimize N_2_O levels in nitrification/denitrification process.

**Table 3.** Nitrogen, phosphorus, and carbon mass balance.

|  | Period-I (g/d) |  |  | Period-II (g/d) |  |  |
| --- | --- | --- | --- | --- | --- | --- |
|  | TN | TP | COD | TN | TP | COD |
| Inf-soluble | 1.13 | 0.25 | 6.17 | 1.13 | 0.108 | 8.91 |
| Eff-soluble | 0.51 | 0.11 | 3.43 | 0.22 | 0.053 | 4.17 |
| Sludge | 0.32 | 0.13 | 1.39 | 0.21 | 0.066 | 2.18 |
| N<sub>2</sub>O | 0.3 | / | / | 0.46 | / | / |
| recovery | 0% | -4% | -22% | -21% | +10% | -25.6 |

Though the P release/uptake content was low in our system, the P content in sludge is around 4∼5% (mg/mg-VSS), within the P content in normal EBPR system treating wastewater and manure wastewater (2∼12%).^10,57,58^ Except biotic phosphorus removal, abiotic phosphorus removal by chemical precipitation also involved in our system and they contributed to 52%-61% P removal in total (Table 3). The chemical precipitation depends on the influent calcium and P concentrations, and the biologically released P concentrations in anaerobic phase. The benefits of integrated partial nitrification/denitrification via N_2_O recovery and phosphorus removal includes a reduced aeration requirement, potential of P recovery, and enhanced energy recovery. Additional testing and research such as N_2_O recovery by membrane-based stripping,^59^ and phosphorus recovery in forms of chemical phosphorus precipitation or poly-P in Ca-rich manure digestate are needed.

## 4. Acknowledgements

We acknowledge Dr. Mi Nguyen for contributing the operational data in Period I. We also acknowledge Dr. Ali Akbari for contributing part of the operational data in Period II. This study was supported by the project Manure Management: Enhanced Nutrient Recovery, joint research between the Atkinson Center for a Sustainable Future (Cornell University) and the Environmental Defense Fund (EDF).

## For Table of Content Only

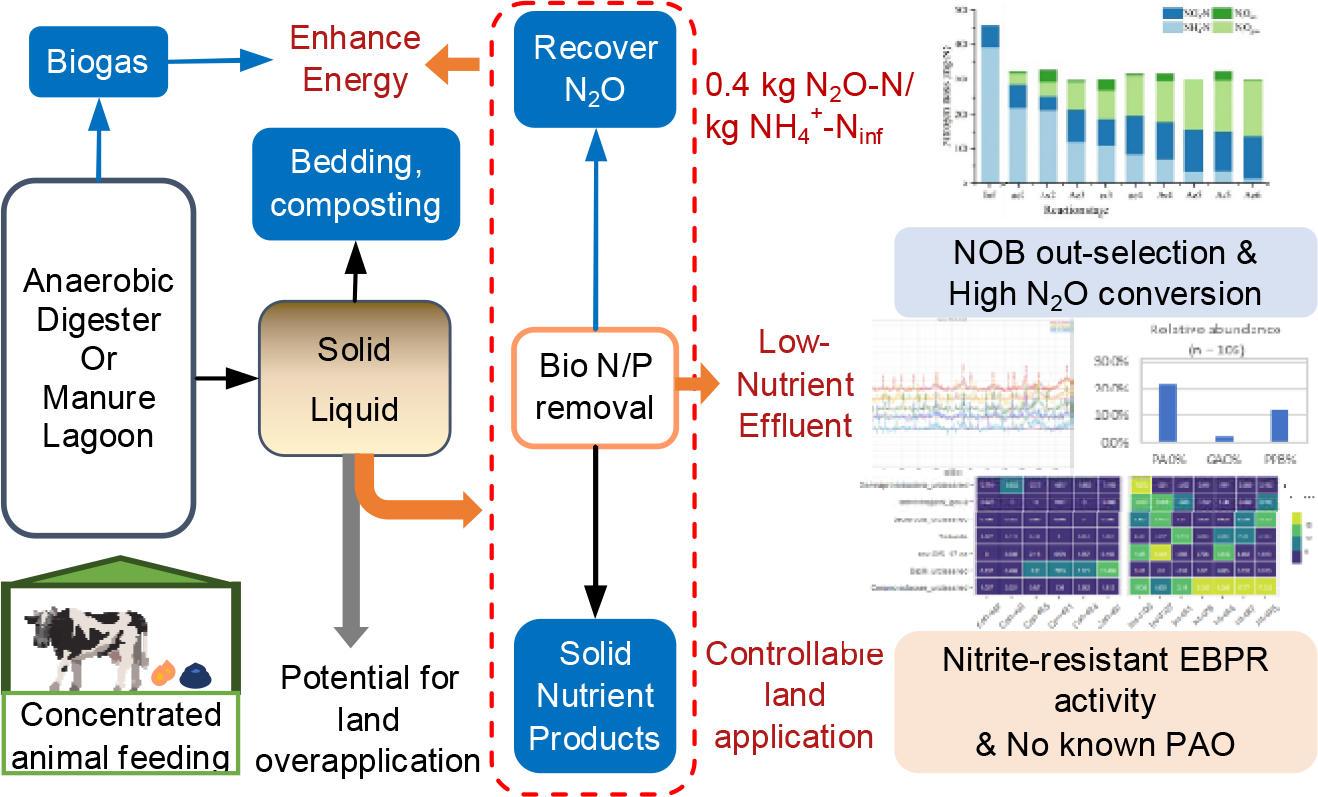

## Notes

### Competing Interest Statement

The authors have declared no competing interest.

