## Supplementary material for "Combined Partial-Nitrification and Phosphorus Removal with the co-Existence of Nitrite-resistant phosphorous accumulating organisms (PAOs) and nitrifiers in the treatment of high-strength manure digestate": Suplemental Information

**Contents**:

Figure:5

Tables:4

Text: 1

Continuous Aeration

DO (mg/L)

|  | DO probe (mg/L) |
| --- | --- |
| 0:00:02 | 3.67081 |
| 0:00:07 | 3.67721 |
| 0:00:12 | 3.65845 |
| 0:00:17 | 3.64487 |
| 0:00:22 | 3.69565 |
| 0:00:27 | 3.71782 |
| 0:00:32 | 3.69215 |
| 0:00:37 | 3.70255 |
| 0:00:42 | 3.70591 |
| 0:00:47 | 3.70368 |
| 0:00:52 | 3.69677 |
| 0:00:57 | 3.67057 |
| 0:01:02 | 3.66689 |
| 0:01:07 | 3.6641 |
| 0:01:12 | 3.70594 |
| 0:01:17 | 3.70396 |
| 0:01:22 | 3.71153 |
| 0:01:27 | 3.69522 |
| 0:01:32 | 3.70447 |
| 0:01:37 | 3.70308 |
| 0:01:42 | 3.69419 |
| 0:01:47 | 3.71238 |
| 0:01:52 | 3.72755 |
| 0:01:57 | 3.71066 |
| 0:02:02 | 3.67258 |
| 0:02:07 | 3.67638 |
| 0:02:12 | 3.69706 |
| 0:02:17 | 3.73445 |
| 0:02:22 | 3.71207 |
| 0:02:27 | 3.6787 |
| 0:02:32 | 3.68411 |
| 0:02:37 | 3.68878 |
| 0:02:42 | 3.6916 |
| 0:02:47 | 3.66837 |
| 0:02:52 | 3.66613 |
| 0:02:57 | 3.67012 |
| 0:03:02 | 3.67686 |
| 0:03:07 | 3.68325 |
| 0:03:12 | 3.69318 |
| 0:03:17 | 3.71663 |
| 0:03:22 | 3.70501 |
| 0:03:27 | 3.67239 |
| 0:03:32 | 3.66222 |
| 0:03:37 | 3.65372 |
| 0:03:42 | 3.71943 |
| 0:03:47 | 3.71251 |
| 0:03:52 | 3.68862 |
| 0:03:57 | 3.68802 |
| 0:04:02 | 3.66743 |
| 0:04:07 | 3.66707 |
| 0:04:12 | 3.67987 |
| 0:04:17 | 3.68817 |
| 0:04:22 | 3.67571 |
| 0:04:27 | 3.69884 |
| 0:04:32 | 3.70139 |
| 0:04:37 | 3.69996 |
| 0:04:42 | 3.6947 |
| 0:04:47 | 3.68099 |
| 0:04:52 | 3.68954 |
| 0:04:57 | 3.6719 |
| 0:05:02 | 3.68177 |
| 0:05:07 | 3.73549 |
| 0:05:12 | 3.75478 |
| 0:05:17 | 3.71673 |
| 0:05:22 | 3.70307 |
| 0:05:27 | 3.71934 |
| 0:05:32 | 3.78808 |
| 0:05:37 | 3.7718 |
| 0:05:42 | 3.7671 |
| 0:05:47 | 3.79033 |
| 0:05:52 | 3.76155 |
| 0:05:57 | 3.71837 |
| 0:06:02 | 3.69354 |
| 0:06:07 | 3.7271 |
| 0:06:12 | 3.72812 |
| 0:06:17 | 3.71559 |
| 0:06:22 | 3.73603 |
| 0:06:27 | 3.73731 |
| 0:06:32 | 3.71229 |
| 0:06:37 | 3.70233 |
| 0:06:42 | 3.71664 |
| 0:06:47 | 3.71336 |
| 0:06:52 | 3.73061 |
| 0:06:57 | 3.73171 |
| 0:07:02 | 3.7586 |
| 0:07:07 | 3.71705 |
| 0:07:12 | 3.72727 |
| 0:07:17 | 3.72136 |
| 0:07:22 | 3.75042 |
| 0:07:27 | 3.74696 |
| 0:07:32 | 3.76321 |
| 0:07:37 | 3.73871 |
| 0:07:42 | 3.74211 |
| 0:07:47 | 3.73604 |
| 0:07:52 | 3.73569 |
| 0:07:57 | 3.7249 |
| 0:08:02 | 3.75136 |
| 0:08:07 | 3.82319 |
| 0:08:12 | 3.77932 |
| 0:08:17 | 3.7547 |
| 0:08:22 | 3.73679 |
| 0:08:27 | 3.74434 |
| 0:08:32 | 3.7819 |
| 0:08:37 | 3.7601 |
| 0:08:42 | 3.76369 |
| 0:08:47 | 3.73883 |
| 0:08:52 | 3.73088 |
| 0:08:57 | 3.72707 |
| 0:09:02 | 3.72432 |
| 0:09:07 | 3.77145 |
| 0:09:12 | 3.7737 |
| 0:09:17 | 3.75456 |
| 0:09:22 | 3.79833 |
| 0:09:27 | 3.80779 |
| 0:09:32 | 3.78606 |
| 0:09:37 | 3.78161 |
| 0:09:42 | 3.8403 |
| 0:09:47 | 3.79538 |
| 0:09:52 | 3.75614 |
| 0:09:57 | 3.76782 |
| 0:10:02 | 3.80878 |
| 0:10:07 | 3.82691 |
| 0:10:12 | 3.77962 |
| 0:10:17 | 3.77701 |
| 0:10:22 | 3.77273 |
| 0:10:27 | 3.80502 |
| 0:10:32 | 3.79073 |
| 0:10:37 | 3.77942 |
| 0:10:42 | 3.81179 |
| 0:10:47 | 3.89643 |
| 0:10:52 | 3.85683 |
| 0:10:57 | 3.78879 |
| 0:11:02 | 3.76313 |
| 0:11:07 | 3.75558 |
| 0:11:12 | 3.76401 |
| 0:11:17 | 3.75016 |
| 0:11:22 | 3.75381 |
| 0:11:27 | 3.73628 |
| 0:11:32 | 3.79698 |
| 0:11:37 | 3.7832 |
| 0:11:42 | 3.75838 |
| 0:11:47 | 3.73106 |
| 0:11:52 | 3.74063 |
| 0:11:57 | 3.80864 |
| 0:12:02 | 3.77546 |
| 0:12:07 | 3.82736 |
| 0:12:12 | 3.83177 |
| 0:12:17 | 3.83337 |
| 0:12:22 | 3.79801 |
| 0:12:27 | 3.77237 |
| 0:12:32 | 3.76544 |
| 0:12:37 | 3.76481 |
| 0:12:42 | 3.76249 |
| 0:12:47 | 3.75774 |
| 0:12:52 | 3.75746 |
| 0:12:57 | 3.78674 |
| 0:13:02 | 3.80204 |
| 0:13:07 | 3.80251 |
| 0:13:12 | 3.784 |
| 0:13:17 | 3.81834 |
| 0:13:22 | 3.82003 |
| 0:13:27 | 3.8326 |
| 0:13:32 | 3.84959 |
| 0:13:37 | 3.84984 |
| 0:13:42 | 3.83565 |
| 0:13:47 | 3.81297 |
| 0:13:52 | 3.78474 |
| 0:13:57 | 3.77419 |
| 0:14:02 | 3.80219 |
| 0:14:07 | 3.81162 |
| 0:14:12 | 3.83991 |
| 0:14:17 | 3.87374 |
| 0:14:22 | 3.86999 |
| 0:14:27 | 3.83115 |
| 0:14:32 | 3.78413 |
| 0:14:37 | 3.8388 |
| 0:14:42 | 3.8452 |
| 0:14:47 | 3.80548 |
| 0:14:52 | 3.7979 |
| 0:14:57 | 3.83045 |
| 0:15:02 | 3.83801 |
| 0:15:07 | 3.84578 |
| 0:15:12 | 3.85384 |
| 0:15:17 | 3.8683 |
| 0:15:22 | 3.88712 |
| 0:15:27 | 3.86437 |
| 0:15:32 | 3.85761 |
| 0:15:37 | 3.84077 |
| 0:15:42 | 3.84976 |
| 0:15:47 | 3.85825 |
| 0:15:52 | 3.86277 |
| 0:15:57 | 3.86147 |
| 0:16:02 | 3.88218 |
| 0:16:07 | 3.86281 |
| 0:16:12 | 3.83318 |
| 0:16:17 | 3.81998 |
| 0:16:22 | 3.87265 |
| 0:16:27 | 3.83818 |
| 0:16:32 | 3.81923 |
| 0:16:37 | 3.82408 |

Intermittent Aeration

DO (mg/L)

|  | DO probe (mg/L) |
| --- | --- |
| 0:00:04 | -0.0192623 |
| 0:00:09 | -0.0187 |
| 0:00:14 | -0.01781 |
| 0:00:19 | -0.0193566 |
| 0:00:24 | -0.0185074 |
| 0:00:29 | -0.0185376 |
| 0:00:34 | -0.0192914 |
| 0:00:39 | -0.018844 |
| 0:00:44 | -0.0184247 |
| 0:00:49 | -0.0185677 |
| 0:00:54 | -0.0196037 |
| 0:00:59 | -0.0194675 |
| 0:01:04 | -0.0182953 |
| 0:01:09 | -0.0190979 |
| 0:01:14 | -0.0190512 |
| 0:01:19 | -0.0185142 |
| 0:01:24 | -0.0191115 |
| 0:01:29 | -0.0196951 |
| 0:01:34 | -0.0195667 |
| 0:01:39 | -0.0183907 |
| 0:01:44 | -0.019664 |
| 0:01:49 | -0.0193284 |
| 0:01:54 | -0.0189597 |
| 0:01:59 | -0.0196728 |
| 0:02:04 | -0.0192525 |
| 0:02:09 | -0.019485 |
| 0:02:14 | -0.0196416 |
| 0:02:19 | -0.0196562 |
| 0:02:24 | -0.0192632 |
| 0:02:29 | -0.0206708 |
| 0:02:34 | -0.0195872 |
| 0:02:39 | -0.0198158 |
| 0:02:44 | -0.0195113 |
| 0:02:49 | -0.0200191 |
| 0:02:54 | -0.0191193 |
| 0:02:59 | -0.0199918 |
| 0:03:04 | -0.0202622 |
| 0:03:09 | -0.0196377 |
| 0:03:14 | -0.0210161 |
| 0:03:19 | -0.0199694 |
| 0:03:24 | -0.0208867 |
| 0:03:29 | -0.0202846 |
| 0:03:34 | -0.0206329 |
| 0:03:39 | -0.0205541 |
| 0:03:44 | -0.0205521 |
| 0:03:49 | -0.0199704 |
| 0:03:54 | -0.0199704 |
| 0:03:59 | -0.0203576 |
| 0:04:04 | -0.020486 |
| 0:04:09 | -0.020521 |
| 0:04:14 | -0.0202097 |
| 0:04:19 | -0.0209169 |
| 0:04:24 | -0.0200093 |
| 0:04:29 | -0.019807 |
| 0:04:34 | -0.0206348 |
| 0:04:39 | -0.0208605 |
| 0:04:44 | -0.0206543 |
| 0:04:49 | -0.0202476 |
| 0:04:54 | -0.0209509 |
| 0:04:59 | -0.0200181 |
| 0:05:04 | -0.0207924 |
| 0:05:09 | -0.0213332 |
| 0:05:14 | -0.020199 |
| 0:05:19 | -0.0204801 |
| 0:05:24 | -0.0203041 |
| 0:05:29 | -0.0207321 |
| 0:05:34 | -0.0210648 |
| 0:05:39 | -0.0202301 |
| 0:05:44 | -0.020592 |
| 0:05:49 | -0.0211241 |
| 0:05:54 | -0.0206727 |
| 0:05:59 | -0.0208031 |
| 0:06:04 | -0.0207943 |
| 0:06:09 | -0.0207749 |
| 0:06:14 | -0.0204967 |
| 0:06:19 | -0.0214354 |
| 0:06:24 | -0.02102 |
| 0:06:29 | -0.0203488 |
| 0:06:34 | -0.0206221 |
| 0:06:39 | -0.0209325 |
| 0:06:44 | -0.0216309 |
| 0:06:49 | -0.0212904 |
| 0:06:54 | -0.0212856 |
| 0:06:59 | -0.0208517 |
| 0:07:04 | -0.0210998 |
| 0:07:09 | -0.0205346 |
| 0:07:14 | -0.0208313 |
| 0:07:19 | -0.0217379 |
| 0:07:24 | -0.0220287 |
| 0:07:29 | -0.0208751 |
| 0:07:34 | -0.0216688 |
| 0:07:39 | -0.0216834 |
| 0:07:44 | -0.0212184 |
| 0:07:49 | -0.0210823 |
| 0:07:54 | -0.0212729 |
| 0:07:59 | -0.0213935 |
| 0:08:04 | -0.0210949 |
| 0:08:09 | -0.0211095 |
| 0:08:14 | -0.0210862 |
| 0:08:19 | -0.020591 |
| 0:08:24 | -0.0218828 |
| 0:08:29 | -0.0218478 |
| 0:08:34 | -0.0215346 |
| 0:08:39 | -0.0211737 |
| 0:08:44 | -0.0213634 |
| 0:08:49 | -0.021769 |
| 0:08:54 | -0.0215054 |
| 0:08:59 | -0.0213877 |
| 0:09:04 | -0.0222914 |
| 0:09:09 | -0.0211601 |
| 0:09:14 | -0.0219723 |
| 0:09:19 | -0.0207661 |
| 0:09:24 | -0.0225355 |
| 0:09:29 | -0.0219188 |
| 0:09:34 | -0.0216241 |
| 0:09:39 | -0.0215239 |
| 0:09:44 | -0.0221688 |
| 0:09:49 | -0.022447 |
| 0:09:54 | -0.0213099 |
| 0:09:59 | -0.0219733 |
| 0:10:04 | -0.021664 |
| 0:10:09 | -0.0208867 |
| 0:10:14 | -0.0222038 |
| 0:10:19 | -0.0222846 |
| 0:10:24 | -0.0215852 |
| 0:10:29 | -0.0222729 |
| 0:10:34 | -0.021664 |
| 0:10:39 | -0.0214762 |
| 0:10:44 | -0.0220248 |
| 0:10:49 | -0.0219023 |
| 0:10:54 | -0.0225521 |
| 0:10:59 | -0.0221698 |
| 0:11:04 | -0.021913 |
| 0:11:09 | -0.0222087 |
| 0:11:14 | -0.0219276 |
| 0:11:19 | -0.0223284 |
| 0:11:24 | -0.0216581 |
| 0:11:29 | -0.0220861 |
| 0:11:34 | -0.0219538 |
| 0:11:39 | -0.0227106 |
| 0:11:44 | -0.022663 |
| 0:11:49 | -0.0228371 |
| 0:11:54 | -0.0228098 |
| 0:11:59 | -0.0230968 |
| 0:12:04 | -0.0236026 |
| 0:12:09 | -0.0236814 |
| 0:12:14 | -0.0231211 |
| 0:12:19 | -0.0237203 |
| 0:12:24 | -0.0241542 |
| 0:12:29 | -0.0234324 |
| 0:12:34 | -0.02359 |
| 0:12:39 | -0.0249693 |
| 0:12:44 | -0.0249509 |
| 0:12:49 | -0.02384 |
| 0:12:54 | -0.0238001 |
| 0:12:59 | -0.0232622 |
| 0:13:04 | -0.0239275 |
| 0:13:09 | -0.0247262 |
| 0:13:14 | -0.0240452 |
| 0:13:19 | -0.0241172 |
| 0:13:24 | -0.0238478 |
| 0:13:29 | -0.0245783 |
| 0:13:34 | -0.0238361 |
| 0:13:39 | -0.0246795 |
| 0:13:44 | -0.024269 |
| 0:13:49 | -0.0241989 |
| 0:13:54 | -0.0240958 |
| 0:13:59 | -0.0242262 |
| 0:14:04 | -0.0240822 |
| 0:14:09 | -0.0230813 |
| 0:14:14 | -0.0245404 |
| 0:14:19 | -0.0238711 |
| 0:14:24 | -0.0233595 |
| 0:14:29 | -0.0239246 |
| 0:14:34 | -0.0239519 |
| 0:14:39 | -0.023912 |
| 0:14:44 | -0.024481 |
| 0:14:49 | -0.0234168 |
| 0:14:54 | -0.0235005 |
| 0:14:59 | -0.0239849 |
| 0:15:04 | -0.0244304 |
| 0:15:09 | -0.0247038 |
| 0:15:14 | -0.0237165 |
| 0:15:19 | -0.0232252 |
| 0:15:24 | -0.0242252 |
| 0:15:29 | -0.0240637 |
| 0:15:34 | -0.0240666 |
| 0:15:39 | -0.0238059 |
| 0:15:44 | -0.0240131 |
| 0:15:49 | -0.0246892 |
| 0:15:54 | -0.0249178 |
| 0:15:59 | -0.0237067 |
| 0:16:04 | -0.0234947 |
| 0:16:09 | 0.00148149 |
| 0:16:14 | 0.0545164 |
| 0:16:19 | 0.134471 |
| 0:16:24 | 0.227206 |
| 0:16:29 | 0.310943 |
| 0:16:34 | 0.395881 |
| 0:16:39 | 0.490043 |

Figure S1 Aeration profiles from (A) Period I (An/Aer) system and (B) Period II (An/Ax/Aer) system

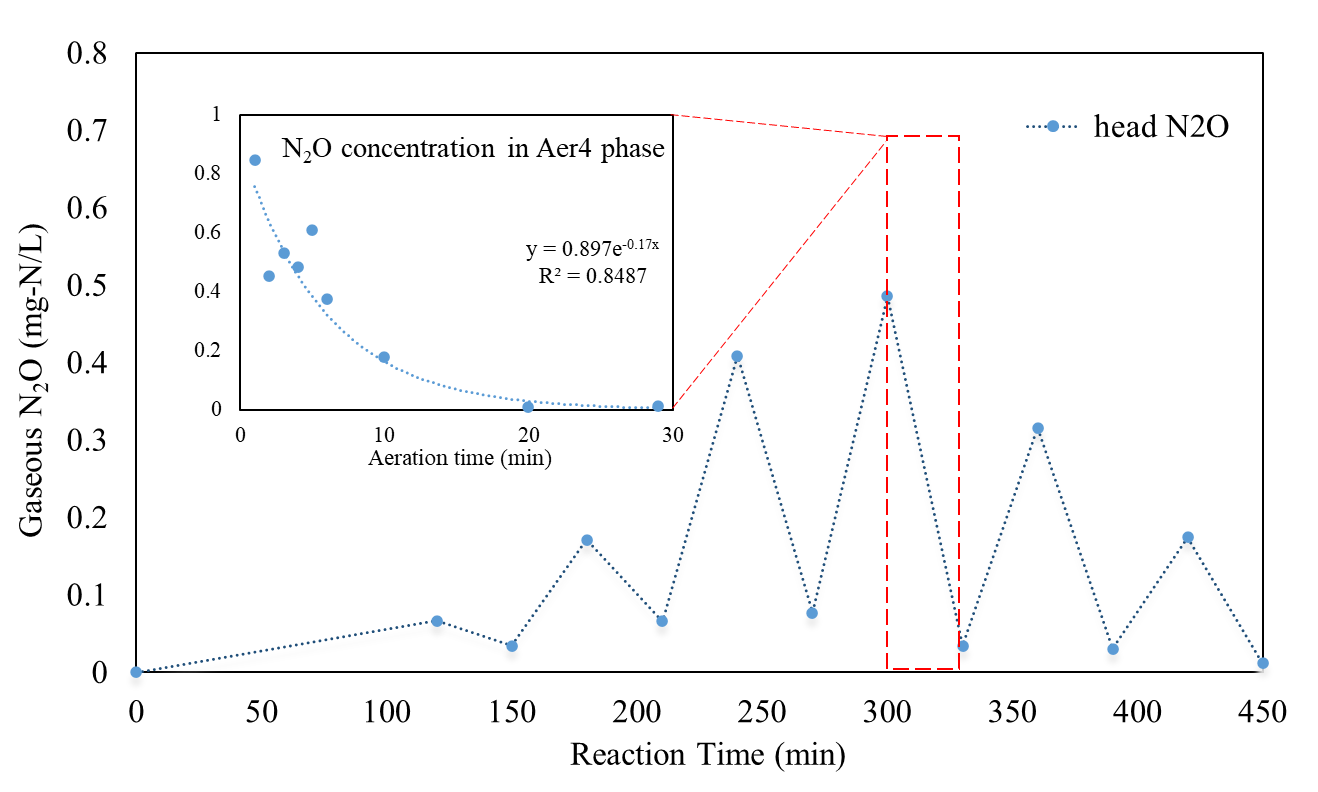

Figure S2 Headspace N_2_O concentration variation during Aeration phase in Period II;

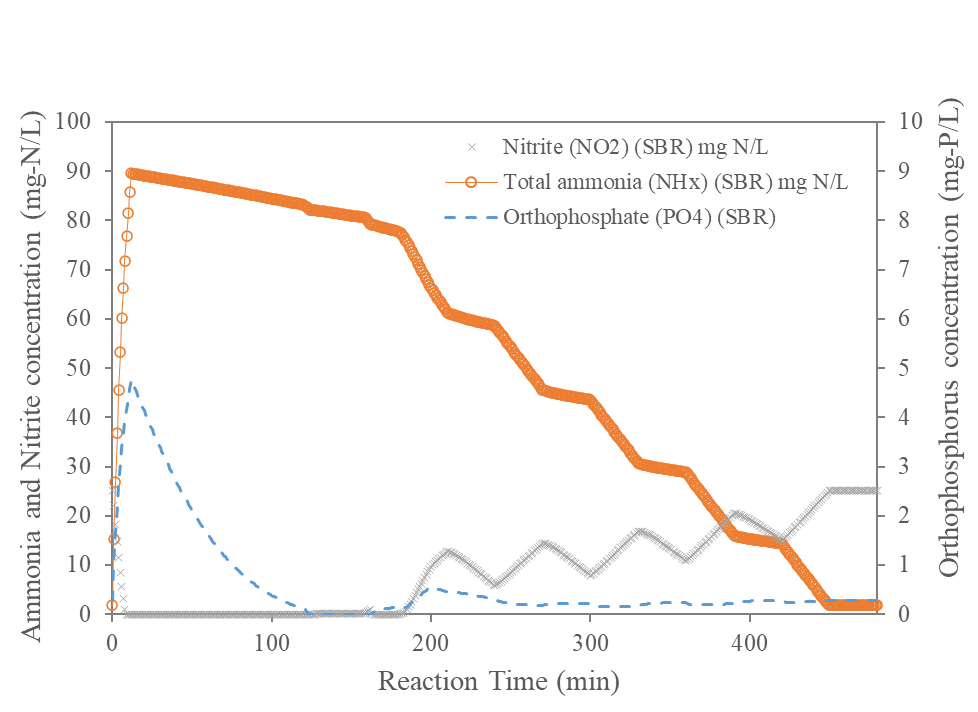

Figure S3 Ammonia, nitrite and orthophosphate concentration in a single cycle modelled by SUMO;

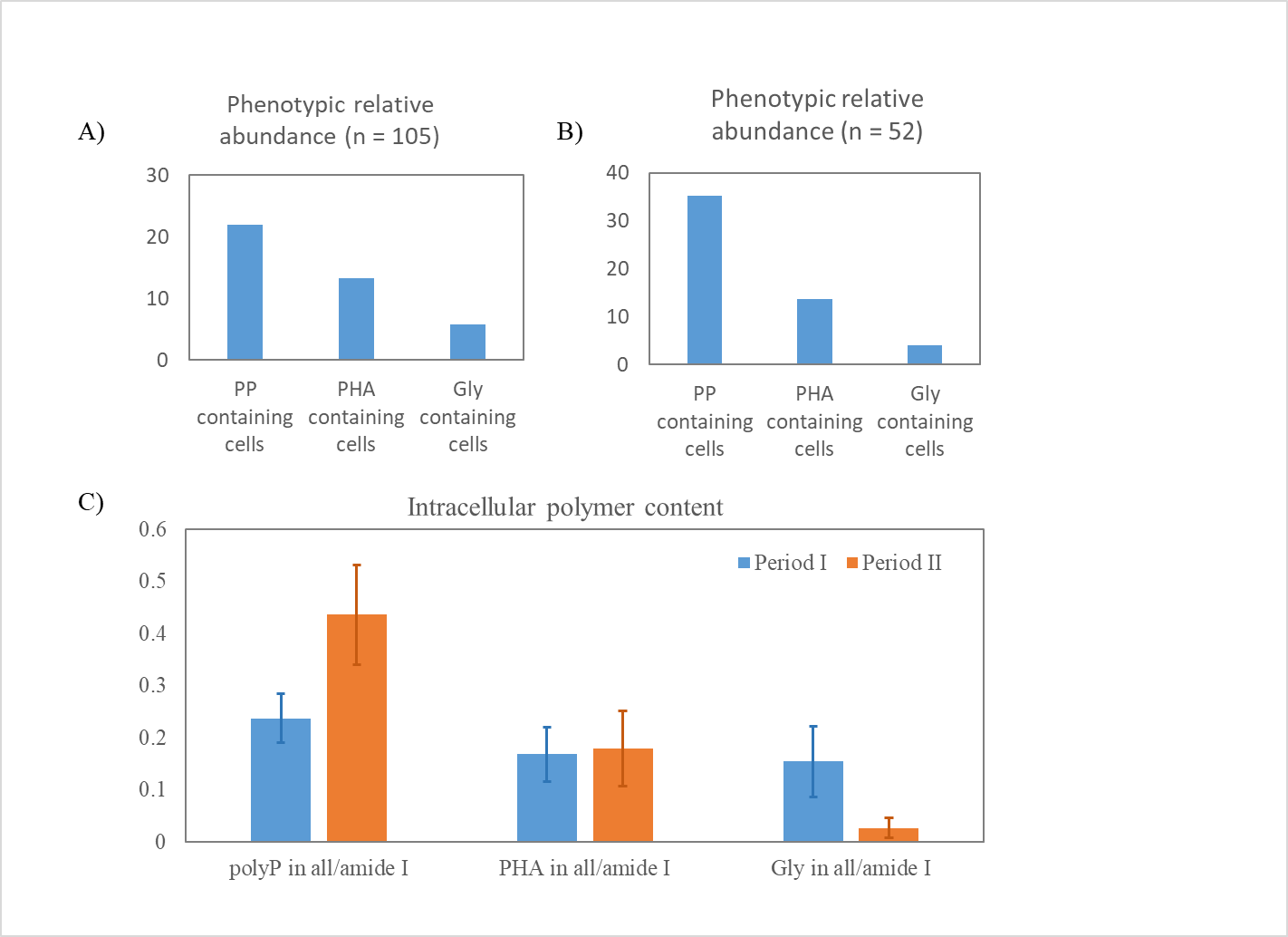

Figure S4 Relative abundance of SCRS featuring with poly-P, PHA and/or glycogen in (A) period I (n = 105) and (B) period II; C) Raman intensity of poly-P, PHA and glycogen normalized by the intensity of amide I within the cell.

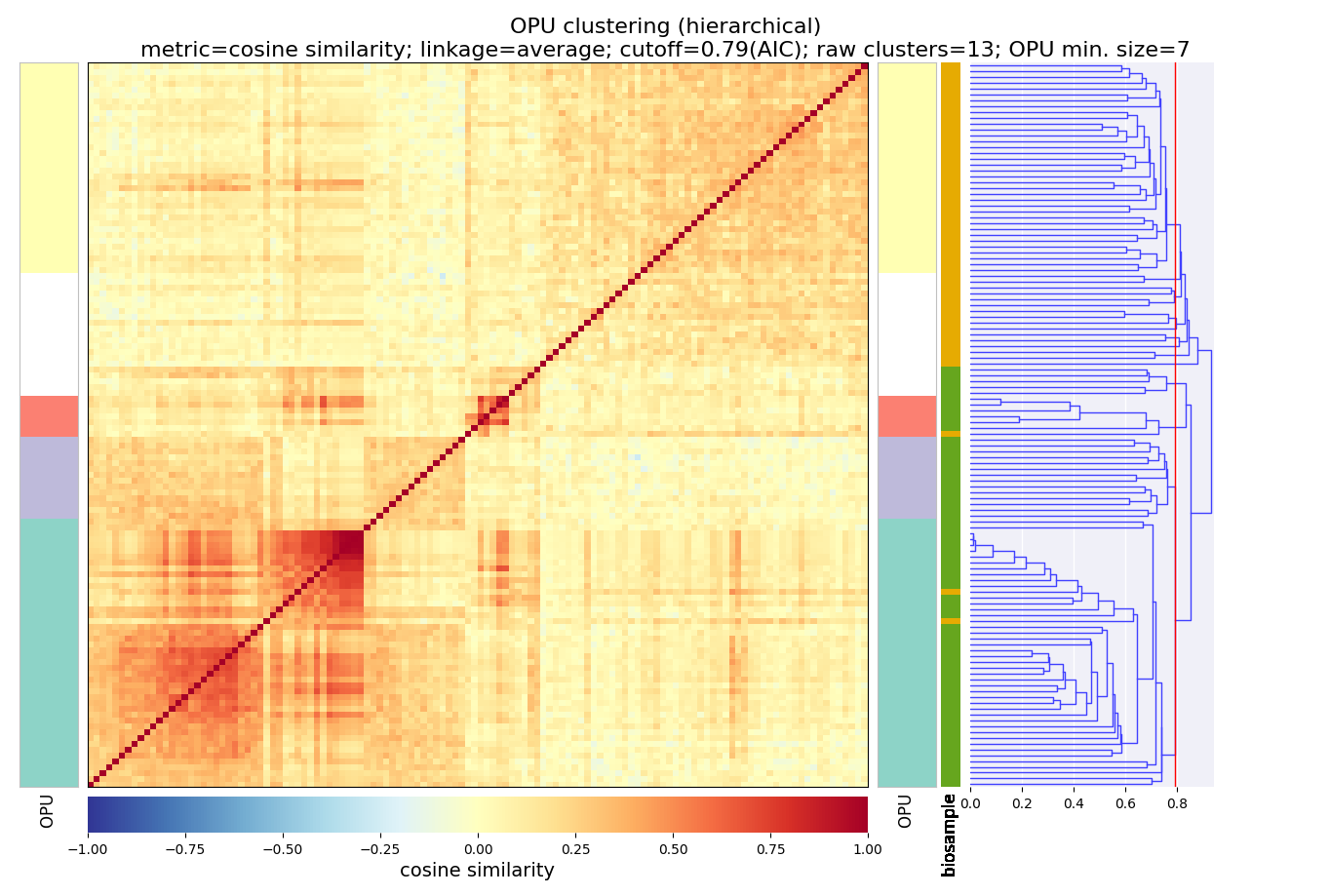

**OPU1**

**OPU2**

**OPU3**

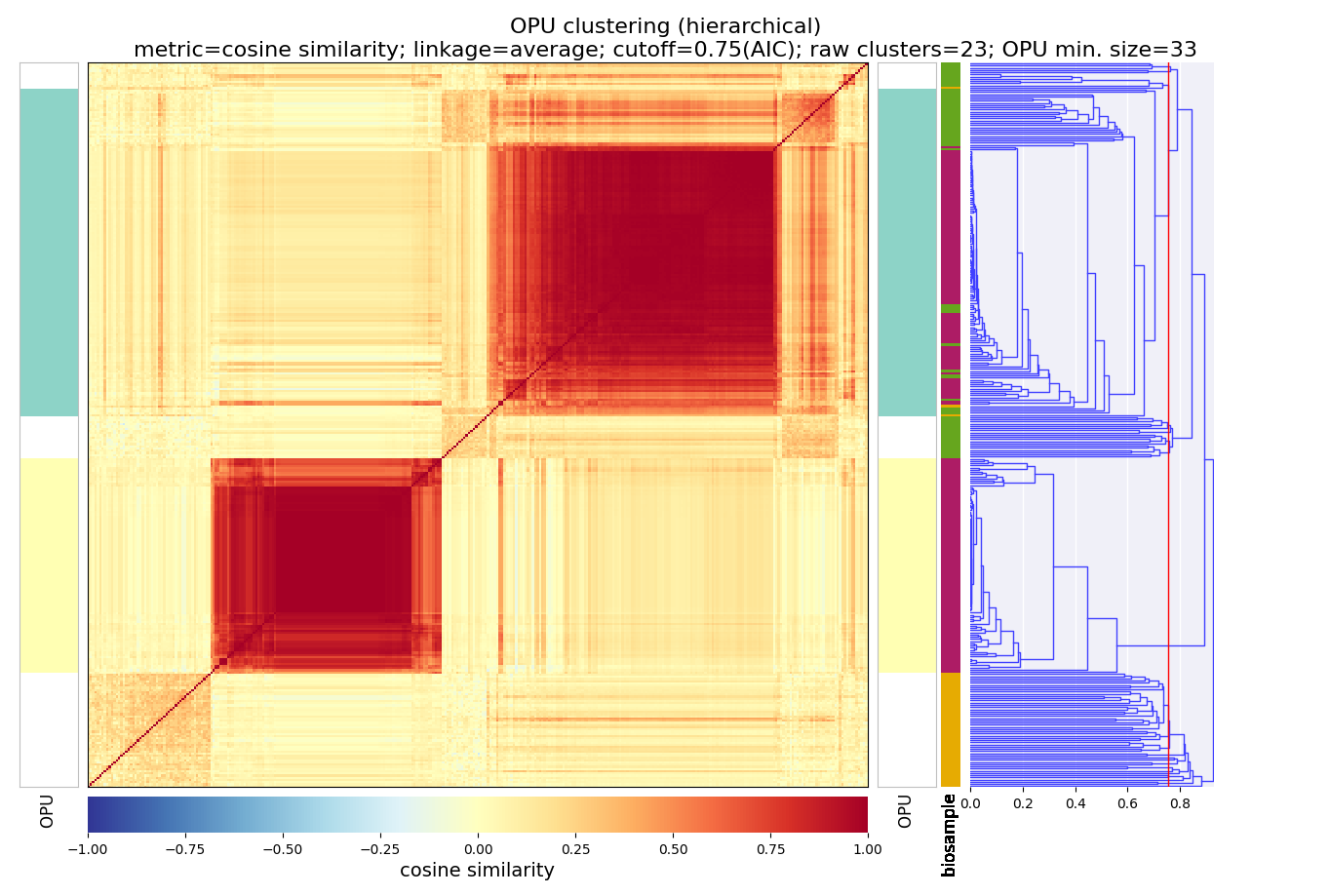

A)

C)

OPU clustering indicate that the SCRS from OPU1 in Figure 3A (biosample color: green) is close to Comamonadaceae labelled SCRS (biosample color: red)

Figure S5 (A) The hierarchical clustering analysis (HCA) of single-cell Raman spectra (SCRS) in period I (biosample color: yellow) and period II (biosample color: green); and (B) The HCA analysis of SCRS in the present study with Comamonadaceae labelled SCRS (biosample color: red).

**Table S1 Literature summary of characteristics of manure wastewaters**

| Wastewater type | COD  (mg/L) | TP  (mg P/L) | NH_4_^+^-N  (mg N/L) | Reference |
| --- | --- | --- | --- | --- |
| Digested dairy manure | 942 ± 12 | 34 ± 2 | 136 ± 8 | ^1^ |
| 20X Diluted digested dairy manure | 1188 | 12.5 | 111.6 | ^2^ |
| 10X Diluted digested dairy manure | 1032 | 11.2 | 155.4 | ^3^ |
| 5X Diluted digested dairy manure | 8412-9751 | 30-90 | 96.2-103.8 | ^4^ |
| 5X Diluted digested dairy manure | NA | 41.9±16.7 | 86.8±24.1 | ^5^ |
| 10X Diluted dairy manure | 2124 ± 472 | 51.1±23.0 | 103 ± 23.4 | ^6^ |
| 10X Diluted digested dairy manure | 2583 ± 521 | 75-180 | NA | ^7^ |
| Raw liquid swine manure | 20666 ± 13197 | 494 ± 228 | 1251 ± 616 | ^8^ |
| Swine manure digestate | 1660 | 82 | 520 | ^9^ |

NA: not available.

Table S2 Summary of medium compositions and nitrogen removal activities in batch tests

| # | Activity test | Acetate | NO_3_^−^-N | NO_2_^−^-N | NH_4_^+^-N | DO | N removal activity |
| --- | --- | --- | --- | --- | --- | --- | --- |
|  |  | mg/L | mg-N/L | mg-N/L | mg-N/L | mg/L | mg-N/g-VSS/h |
| 1 | Anaerobic  Heterotrophic Denitrification | 80 | 20 | - | - | - | 2.94 ± 0.09 |
| 2 | Anaerobic  Heterotrophic Denitritation | 80 | - | 20 | - | - | 2.77 ± 0.13 |
| 3 | Aerobic  Heterotrophic Denitrification | 80 | 20 | - | - | 2.0 | 0 |
| 4 | Endogenous  Denitrification | - | 20 | - | - | - | 0.20 ± 0.01 |
| 5 | Anammox | - | - | 20 | 15 | - | 1.32 ± 0.03 |

| Table S3 Manure digestate constitutes modeled in Visual MINTEQ | | | | |  |
| --- | --- | --- | --- | --- | --- |
| Name | Unit | T_0_ | T_120_ | Eff. | Quantification Method |
| pH | 1 | 7.00 | 7.20 | 6.66 | HACH portable pH probe |
| ORP | mV | -302.00 | -148 | 127 | HACH portable ORP probe |
| VFA (as acetate) | mg-C/L | 153 | 100 | 96 | HACH TNT872 |
| NH_4_^+^-N | mg-N/L | 70.0 | 68.0 | 3.8 | IC |
| NO_2_^−^-N | mg-N/L | 15.0 | 2.4 | 31.1 | IC |
| TP | mg-P/L | 14.0 | 23.6 | 9.6 | ICP-MS |
| SO_4_^2−^ | mg/L | 1.9 | 4.6 | 7.2 | IC |
| Cl^−^ | mg/L | 369 | 348 | 326 | IC |
| Ca | mg/L | 56 | 53.5 | 51 | ICP-MS |
| Mg | mg/L | 23.1 | 21.0 | 19.0 | ICP-MS |
| Al | mg/L | 0.3 | 0.2 | 0.1 | ICP-MS |
| Parameters unchanged | | | | |  |
| Fe | mg/L | 1.4 | | | ICP-MS |
| Alkalinity | mg CaCO3/L | 1960 | | | HACH Alkalinity test |
| DOC | mg/L | 650 | | |  |

| Table S4 Phosphate precipitation calculation by Minteq at the end of anaerobic phase (Unit: mol/L) | | | | | | | | |
| --- | --- | --- | --- | --- | --- | --- | --- | --- |
| Component | Dissolved inorganic | Bound to DOM | Total dissolved | % dissolved | Total sorbed | % sorbed | Total precipitated | % precipitated |
| >SOH(1) | 0 | 0 | 0 | 0 | 4.2504E-20 | 100 | 0 | 0 |
| Acetate-1 | 0.0016936 | 0 | 0.0016936 | 100 | 0 | 0 | 0 | 0 |
| Al+3 | 7.6425E-12 | 7.4125E-06 | 7.4125E-06 | 100 | 0 | 0 | 0 | 0 |
| Ca+2 | 0.000045607 | 0.00049086 | 0.00053647 | 34.626 | 6.4419E-29 | 0 | 0.0010128 | 65.374 |
| Cl-1 | 0.0049076 | 3.521E-07 | 0.0049079 | 100 | 0 | 0 | 0 | 0 |
| CO3-2 | 0.023399 | 3.9348E-06 | 0.023403 | 100 | 0 | 0 | 0 | 0 |
| Fe+2 | 0 | 7.0613E-09 | 7.0613E-09 | 100 | 5.0935E-41 | 0 | 0 | 0 |
| Fe+3 | 0 | 0.000025061 | 0.000025061 | 100 | 0 | 0 | 0 | 0 |
| H+1 | 0.026348 | -0.0031871 | 0.023161 | 100 | 6.4923E-21 | 0 | 0 | 0 |
| HFA1-(6)(aq) | 0 | 0 | 0.0063063 | 100 | 0 | 0 | 0 | 0 |
| HFA2-(6)(aq) | 0 | 0 | 0.0019949 | 100 | 0 | 0 | 0 | 0 |
| K+1 | 0.005323 | 0.0010705 | 0.0063935 | 100 | 0 | 0 | 0 | 0 |
| Mg+2 | 0.00014222 | 0.0016265 | 0.0017687 | 100 | 0 | 0 | 0 | 0 |
| NH4+1 | 0.0040435 | 0.00081117 | 0.0048547 | 100 | 0 | 0 | 0 | 0 |
| NO2-1 | 0.00014279 | 0 | 0.00014279 | 100 | 0 | 0 | 0 | 0 |
| PO4-3 | 0.00018561 | 0 | 0.00018561 | 23.397 | 1.2441E-21 | 0 | 0.0006077 | 76.603 |
| SO4-2 | 0.000047365 | 6.0952E-11 | 0.000047365 | 100 | 5.6705E-22 | 0 | 0 | 0 |

**Text S1 Raman spectra data preprocessing methods**

Load the data into software LabSpec 6. Split the array to get the spectrum of each sampling points. Combine the data of ~10 background points with adjusting overlapping intensities to produce the average background spectrum. Screen the data for removing cosmic ray/spike, outlier and poor-quality spectra. Conduct a smoothing, filtering and correction algorithm to filter high-frequency noise in spectra (Size 3, Degree 4, Polynomial type, Correction 1). Conduct a baseline correction algorithm to account for fluorescence interference and subtract averaged background from sample spectrum (Polynomial type, Degree 10, Max points 256, Noise points 64). Find the peaks of the processing spectrum (Amplitude 5%, Size 20 pix). Identify the single-cell spectrum by removing the spectrum with the intensity of either 995-1010 cm^-1^ (bacteria’s phenylalanine) or 1640-1690 cm^-1^ (bacteria’s amide I) as 0.

The statistical sufficiency of the sampling size was evaluated with radial based function kernel and Eigen-decomposition method which showed that the 90% line converges was at around 130 single-cell Raman spectra and the sample size larger than 40 cells has been proved to be sufficient for assessing the Raman-based phenotypic diversity.^10^ A total of 105 and 65 Raman spectra of single cells were obtained finally at each sampling time which was far enough to do the subsequent analysis. The presence of PHA in single cells was identified by the signature peaks in the range of 1715-1740 cm^−1^ based on previous study.^11^The relative intensity of PHA for single cell was normalized by the intensity of amide I vibration at 1640-1690 cm^−1^. The hierarchical clustering analysis (HCA) was applied on all of the single-cell Raman spectra from activated sludge samples to obtain phenotypic profiles based on operational phenotypic units (OPUs) according to (Li et al., 2018). The cosine similarity (√(2-2r), r-correlation efficient) was used to measure metrics between two samples, average linkage was applied to quantify dissimilarities between two clusters and the cutoff threshold for OPUs was set with Akaike information criterion (AIC).
